# B lymphocytes acquire myeloid and autoimmune phenotypes via the downregulation of lymphocyte-specific protein-1

**DOI:** 10.1101/2024.06.28.600734

**Authors:** Naeun Lee, Bong-Ki Hong, Sungyong You, Riri Kwon, Jihoon Kwon, Eunbyeol Choi, Kang-Gu Lee, Yu-Mi Kim, Yingjin Li, Jayhyun Kim, Young-Jun Park, Yeonseok Chung, Sin-Hyeog Im, Laurent Sabbagh, Chul-Soo Cho, Wan-Uk Kim

## Abstract

Actin-binding proteins (ABPs) have been established as important mediators of immune homeostasis, but their effects on lymphocytes are poorly understood. Here, we demonstrated that LSP1, an ABP, is a master regulator for innate immune responses in B lymphocytes. *Lsp1* deficiency in B cells upregulated the expression of myeloid genes, including CD11b, CD11c, and myeloperoxidase, and bestowed myeloid morphology. Strikingly, *Lsp1*-deficient B cells exhibited dual functions, namely, strong phagocytic activity and high antibody (Ab) production, like ‘chimera’. The PKCβ-CEBPβ pathway was found to be required for such functional chimerism. Moreover, *Lsp1* deficiency induced the myeloid B cell phenotype and autoantibody production in B cells and consequently accelerated the progression of experimental lupus in mice. These changes were abrogated by retinoic acid, which upregulated LSP1 expression. In lupus patients, LSP1 expression in B cells was downregulated and inversely correlated with myeloperoxidase (MPO) expression. Overall, this study reveals a new role of the ABP LSP1 in B lymphocytes and emphasizes its critical involvement in promoting autoimmune responses, particularly by generating functionally chimeric B cells.

## Introduction

The adaptive immune system is a specific and acquired immune system found in vertebrates that is mainly composed of two specialized immune cell types, B and T lymphocytes. Over the last decade, a subset of myeloid marker CD11b^+^, CD11c^+^, and/or T-bet^+^ B cells, defined as age/autoimmune-associated B cells (ABCs), has been the focus of increasing interest^1^. These cells are a relatively small B cell subpopulation but have unique immune senescence features that contribute to inflammaging^1^. Moreover, they suppress B lymphopoiesis while generating a distinct antibody (Ab) repertoire with high reactivity to self-antigens in proinflammatory microenvironments, leading to the generation of diverse autoantibodies^2^. Therefore, the abnormal expansion of ABCs has been suggested to play pivotal roles in the development of immune senescence and autoimmune diseases, such as systemic lupus erythematosus (SLE)^3^. However, it remains elusive which regulator(s) or pathway(s) is necessary for the generation of ABCs.

Aging and the proinflammatory milieu have been described to enhance myelopoiesis and attenuate lymphopoiesis^4^. Interestingly, even after hematopoiesis is completed, the conversion of lymphocytes into myeloid cells can occur through the process of lineage conversion, which requires the reprogramming of distinct transcription factors that define cell lineages through chromatin remodelling^5^. A growing body of evidence has suggested that actin cytoskeletal remodeling, in which diverse actin-binding proteins (ABPs) participate, is implicated in the lineage conversion of immune cells^6^. Traditionally, ABPs have been considered key elements that control cellular motility, such as cell migration and chemotaxis. Recent studies have demonstrated that ABPs also regulate other cellular functions, including phagocytosis, cell morphological changes, epithelial–mesenchymal transition, and protein secretion^6,7^. However, it is still unknown which ABPs regulate the myeloidization of lymphocytes and the generation of ABCs under aging and proinflammatory conditions.

Lymphocyte-specific protein 1 (LSP1) is an ABP that negatively regulates the migration of neutrophils and T lymphocytes^8,9,10^. It has Ca^2+-^binding and F-actin-binding domains and functions as a signal regulator that directly binds to protein kinase C (PKC) in B cells^11^. Our group demonstrated that LSP1 expression is downregulated in the lymphocytes of rheumatoid arthritis patients^10^. *Lsp1* deficiency accelerates chronic arthritis by facilitating T-cell migration to inflamed joints and promoting autoantibody production in mice, suggesting its pathogenic role in autoimmune diseases. Despite earlier findings^8–11^, the role of LSP1 in autoimmune diseases, particularly in the generation of ABCs, has yet to be determined.

Here, global interactome analysis revealed that LSP1 was identified as the top-ranked ABP related to innate immunity, despite being expressed in lymphocytes, which are representative of adaptive immune cells. Based on these findings, we propose the intriguing hypothesis that LSP1, as an ABP, could modulate the innate function of lymphocytes, promoting ABC generation. Strikingly, *Lsp1*-deficient B cells exhibited dual functions, namely, strong phagocytic activity and high Ab production, like ‘chimeric’ cells. The PKCβ-CEBPβ pathway was required for such functional chimerism. Moreover, *Lsp1* deficiency accelerated the progression of experimental lupus in mice, promoting the acquisition of myeloid B cells, the generation of ABCs, and the production of autoantibodies. Notably, the LSP1 signature was increased in SLE patients and correlated with disease activity, the type I IFN signature, and myeloperoxidase (MPO) expression in B cells. Overall, our study identified a novel subtype of LSP1-regulated B cells (LRBs) that have both strong phagocytic and high autoantibody-producing abilities, which may play critical roles in the generation of ABCs and the progression of autoimmune diseases, such as SLE.

## Results

### LSP1 is the core ABP related to the immune responses in lymphocytes

Although ABPs are associated with immune responses, including chemotaxis, phagocytosis, and lineage conversion of immune cells, their function in lymphocytes has not been globally analyzed. To explore which ABPs participate in immune responses by lymphocytes, we initially searched for 260 ABPs that are expressed in lymphocytes using a Gene Ontology Molecular Functions (GOMF) analysis and the Human Protein Atlas (**Extended Data Fig. 1a**). As a result, 40 ABPs were found to be expressed in lymphocytes (**Extended Data Fig. 1b**). To predict the involvement of lymphocyte ABPs in immune responses, a protein‒protein interaction (PPI) network analysis of each ABP was conducted utilizing the STRING database, followed by a Gene Ontology Biological Process (GOBP) enrichment analysis (**Extended Data Fig. 1a** and **1c**). Ridgeline plot analysis of the enrichment scores for immune response-related GOBP terms highlighted LSP1, FMNL1, MSN, LASP1, and MPRIP as the top five candidate ABPs (**Fig. 1a** and **Extended Data Fig. 1c**). Notably, LSP1 was observed to be the highest ranked ABP in lymphocytes with distinct association with immune responses, particularly with several aspects of innate immunity (**Fig. 1a, 1b, Extended Data Fig. 1c**, and **Table 1**), including the regulation of cytokine production, NF-κB signaling, and TNF production (**Fig. 1b**). LSP1 was still ranked high even in macrophages and hematopoietic cells, but other ABPs, including NOD2, GBP1, and GCSAM, exhibited a higher enrichment score and a greater percentage of immune response-related GOBP (**Extended Data Table 1**).

**Fig. 1.**
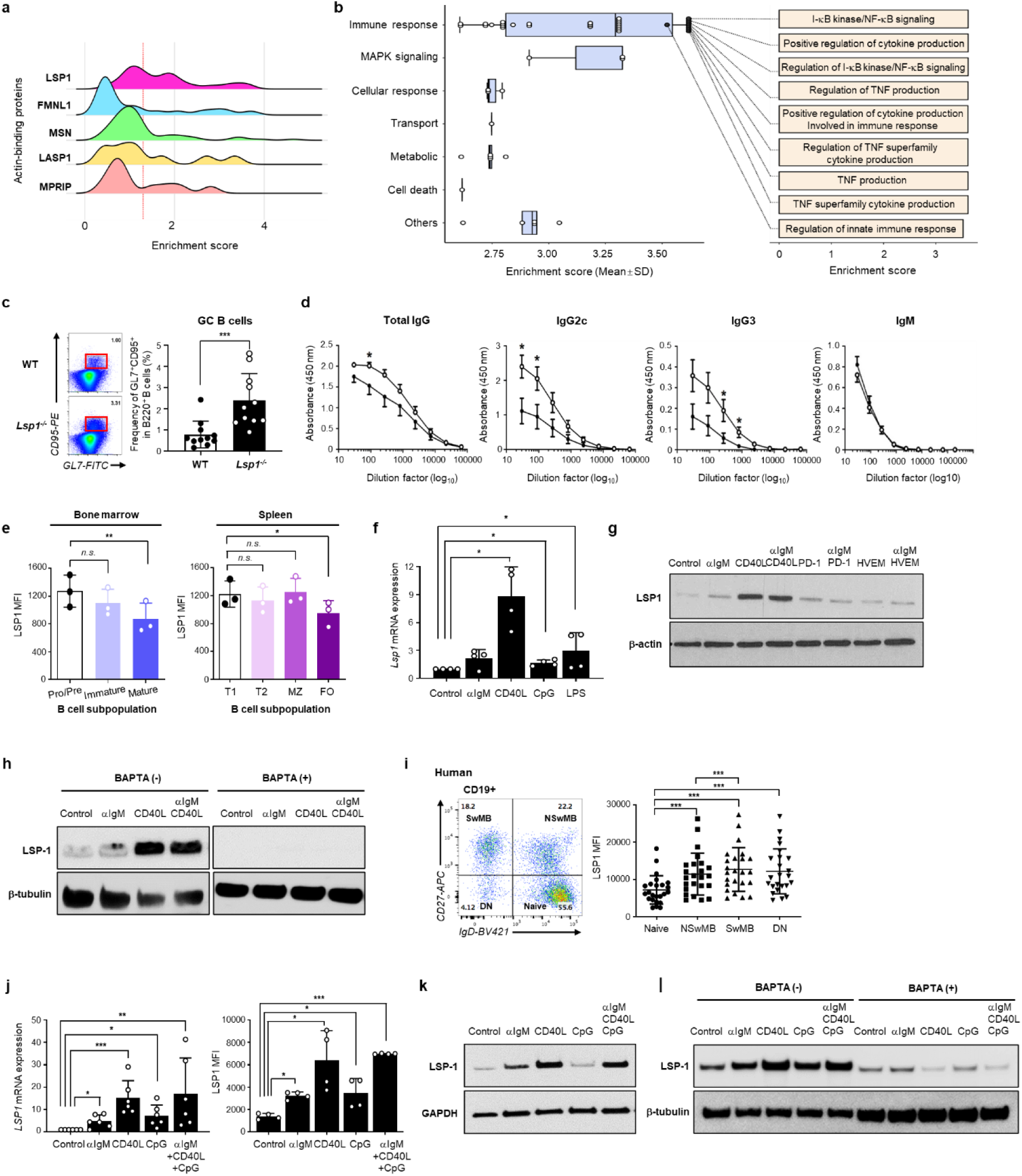
LSP1, a core ABP related to innate immunity in lymphocytes, is expressed throughout the maturation, development, and activation of B cells. **a,** Ridgeline plot showing the enrichment scores of the GOBP terms related to immune responses associated with lymphocyte ABPs, which were determined based on protein-protein interaction (PPI) database. The top five ABPs are presented. b, Enrichment scores of GOBPs determined using the gene sets of the first neighbors of LSP1. The right panel lists the top 10 out of the 33 enriched GOBPs related to immune responses. c, WT and *Lsp1^-/-^* mice were subcutaneously immunized with 100 μg of KLH emulsified in CFA. The percentage of GC B cells (GL7^+^CD95^+^ B220^+^ cells; highlighted in red rectangles) in the draining lymph nodes of mice was analyzed on day 7 (n=11 per group). Representative pseudocolor plots are shown on the left. d, Total IgG, IgG2c, IgG3, and IgM Ab levels in the sera of WT and *Lsp1^-/-^* mice immunized with KLH. e, LSP1 expression across B-cell developmental stages (n=3) determined by flow cytometry. Mouse B cells were derived from the bone marrow (left panel) and spleen (right panel). MFI=mean fluorescence intensity. f, g, qRT‒PCR (n=4; f) and immunoblot (g) analyses of LSP1 expression in B cells. Mouse splenic B cells were stimulated with αIgM Ab (10 μg/mL), CD40L (0.1 μg/mL), CpG (1 μg/mL), LPS (2 μg/mL), PD-1 (5 μg/mL), or HVEM (5 μg/mL) for 3 days. h, Downregulation of LSP1 expression by a calcium chelator. Splenic B cells were stimulated as in (g) in the absence or presence of 5 μM BAPTA and subjected to immunoblotting. i, LSP1 expression according to the differentiation state of human B cells (n=24), which was determined via flow cytometry (left). j, k, qRT‒PCR analysis (n=4; left panel in j), flow cytometric analysis (n=4; right panel in j), and immunoblot analysis (k) of LSP1 expression in human B cells stimulated with the indicated stimuli for 3 days. l, Downregulation of LSP1 expression in human B cells by BAPTA (5 μM). The data in the bar graphs are presented as the means ± SDs of at least two independent experiments. *P* values were determined by the Mann‒Whitney U test (c), two-way analysis of variance (ANOVA) with multiple comparisons (d), and one-way ANOVA with multiple comparisons (e to j). *n.s.*, not significant; * *P* < 0.05; ** *P* < 0.01; *** *P* < 0.001; **** *P* < 0.001.

The PPI network analysis suggested that LSP1, an ABP, is involved primarily in the activation of lymphocytes. To investigate whether LSP1 affects adaptive immune responses, we generated a keyhole limpet hemocyanin (KLH) immunization model in wild-type (WT) and *Lsp1*^-/-^ mice; immunization with KLH has been used to assess T-cell-dependent B-cell responses^12^. We found that *Lsp1* deficiency failed to influence T cell polarization, such as IFN-γ-producing Th1 cells, IL-17-producing Th17 cells, Foxp3-expressing regulatory T cells, and follicular helper T cells (**Extended Data Fig. 2a**). In contrast, *Lsp1*^-/-^ mice exhibited a higher percentage of germinal center (GC) B cells, which was accompanied by increased serum IgG, IgG2c, and IgG3 levels but no change in serum IgM levels (**Fig. 1c**, **1d,** and **Extended Data Fig. 2b**). Taken together, these *in vivo* data suggest that LSP1 negatively regulates B-cell activation and Ab production without affecting T-cell polarization.

### LSP1 expression changes during B-cell maturation, development, and activation

LSP1 is a Ca^2+-^activated protein (52 kDa) expressed in lymphocytes, neutrophils, and macrophages^13,14^. After B-cell receptor (BCR) ligation, LSP1 transmits an activation signal via a PKCβ-dependent pathway^11^, but the dynamics of LSP1 expression throughout B-cell development and activation remain to be defined. To address this issue, we first examined LSP1 expression levels during B-cell development using flow cytometry. As a result, LSP1 expression levels gradually decreased from pro- to pre-B cells to mature B cells in murine bone marrow (**Fig. 1e**). After maturation, follicular B cells expressed lower levels of LSP1 than transitional (T1, T2) and marginal zone (MZ) B cells in the mouse spleen (**Fig. 1e**), indicating that LSP1 expression levels decrease with B-cell maturation and development. In contrast, in murine splenic B cells, *Lsp1* mRNA expression levels modestly increased in response to B-cell receptor (BCR) crosslinking with anti-IgM Ab (αIgM) or lipopolysaccharide (LPS) and markedly in response to stimulation with CD40 ligand (CD40L), a T-cell-dependent costimulatory molecule (**Fig. 1f**). The changes in LSP1 protein expression in response to CD40 ligation were similar (**Fig. 1g** and **Extended Data Fig. 3**), which confirmed that B-cell activation via CD40 ligation upregulates LSP1 expression. Meanwhile, stimulation with programmed cell death protein 1 (PD-1) and herpesvirus entry mediator (HVEM) failed to affect LSP1 expression (**Fig. 1g** and **Extended Data Fig. 3**). It is well known that BCR and/or CD40L ligation increases intracellular Ca^2+^ levels in B cells^15^. The CD40L-induced upregulation of LSP1 expression was almost completely abrogated by the calcium chelator BAPTA, which suggested that this increase was mediated by the Ca^2+^-signaling pathway (**Fig. 1h**).

Human peripheral B cells can be divided into 4 subtypes, naïve, nonswitching memory (NSwMB), switching memory (SwMB), and CD27-IgD-double negative (DN) B cells, depending on their differentiation status^16^. We found that LSP1 levels were greater in antigen-experienced memory B cells (NSwMB, SwMB, and DN B cells) than in naïve B cells and that LSP1 levels increased in proportion to the extent of differentiation of memory B cells (**Fig. 1i**), suggesting that repeated exposure to antigens is linked to LSP1 upregulation in B cells. In support of this notion, *Lsp1* mRNA and protein expression levels increased in response to BCR crosslinking with αIgM and stimulation with toll-like receptor (TLR) 9 ligand CpG in the peripheral B cells of healthy subjects, and this effect was more pronounced in response to CD40 ligation, as determined by qRT‒PCR, flow cytometry, and western blot analysis (**Fig. 1j** and **1k**), which coincided with the mouse data. The LSP1 increase by αIgM and CD40L was also contingent on Ca^2+^-signaling in human B cells (**Fig. 1l**).

In summary, the data from both human and mouse systems demonstrated that LSP1 expression levels gradually decrease during the maturation and development of B cells but increase after maturation upon activation by BCR and/or CD40 ligation, which appears to be regulated by Ca^2+^ signaling.

### Lsp1-deficient B cells display myeloid gene expression profiles

The bioinformatics analysis results shown in **Fig. 1a** and **1b** raised the interesting question of whether LSP1 regulates the innate function of lymphocytes, which are mostly adaptive immune cells. To answer this question, we performed global gene expression analysis through RNA sequencing. We identified 485, 462, and 636 DEGs (724 up- and 232 downregulated genes) from the comparisons of 1) *Lsp1^-/-^*versus WT, 2) *Lsp1^-/-^*+αIgM versus WT+αIgM, and 3) *Lsp1^-/-^* +LPS versus WT+LPS, respectively (**Extended Data Fig. 4a** and **4b**). We then clustered the genes into cluster 1, which comprised the genes upregulated in the *Lsp1^-/-^* B cells stimulated with media, αIgM, and LPS (n=201); cluster 2 (n=298) and cluster 3 (n=172), which comprised of the genes upregulated (excluding cluster 1) and downregulated by either αIgM or LPS in *Lsp1^-/-^*B cells, respectively (**Fig. 2a**, the left panel). Surprisingly, analysis of the cluster 1 and 2 gene sets revealed that the hematopoietic cell lineage pathway and myeloid cell function-related pathways, including the TLR, TNF, and nucleotide-binding oligomerization domain (NOD)-like receptor (NLR) signaling pathways; osteoclast differentiation; and the phagosome pathway, were significantly enriched in the DEGs (**Extended Data Fig. 4c**). Consistent with these results, many genes belonging to the hematopoietic cell lineage pathway, including *Csf1r, Itgam (Cd11b), Cd14, Cd68, Csf3r (Cd114), and Cd177*, were molecular markers for lineage differentiation into macrophages and neutrophils (Fig. 2a, right panel)^17^. The genes in clusters 1 and 2 were also categorized into ligands, receptors, kinases, and transcription factors, which are shown in **Extended Data Fig. 4d**. Consistent with the RNA sequencing data, qPCR confirmed the upregulated expression of myeloid genes, including *Mpo, Cd11b, Cd11c, Tlr7*, *Irf7*, *Tbx21*, *Cybb*, *C3*, and *Il1b,* in *Lsp1^-/-^* B cells (**Fig. 2b**). Meanwhile, the expression levels of genes related to B-cell differentiation into Ab-producing cells, such as *Aicda*, *Prdm1*, and *Irf4*, did not differ between WT and *Lsp1*^-/-^ B cells (**Fig. 2b** and **Extended Data Fig. 5a**).

**Fig. 2.**
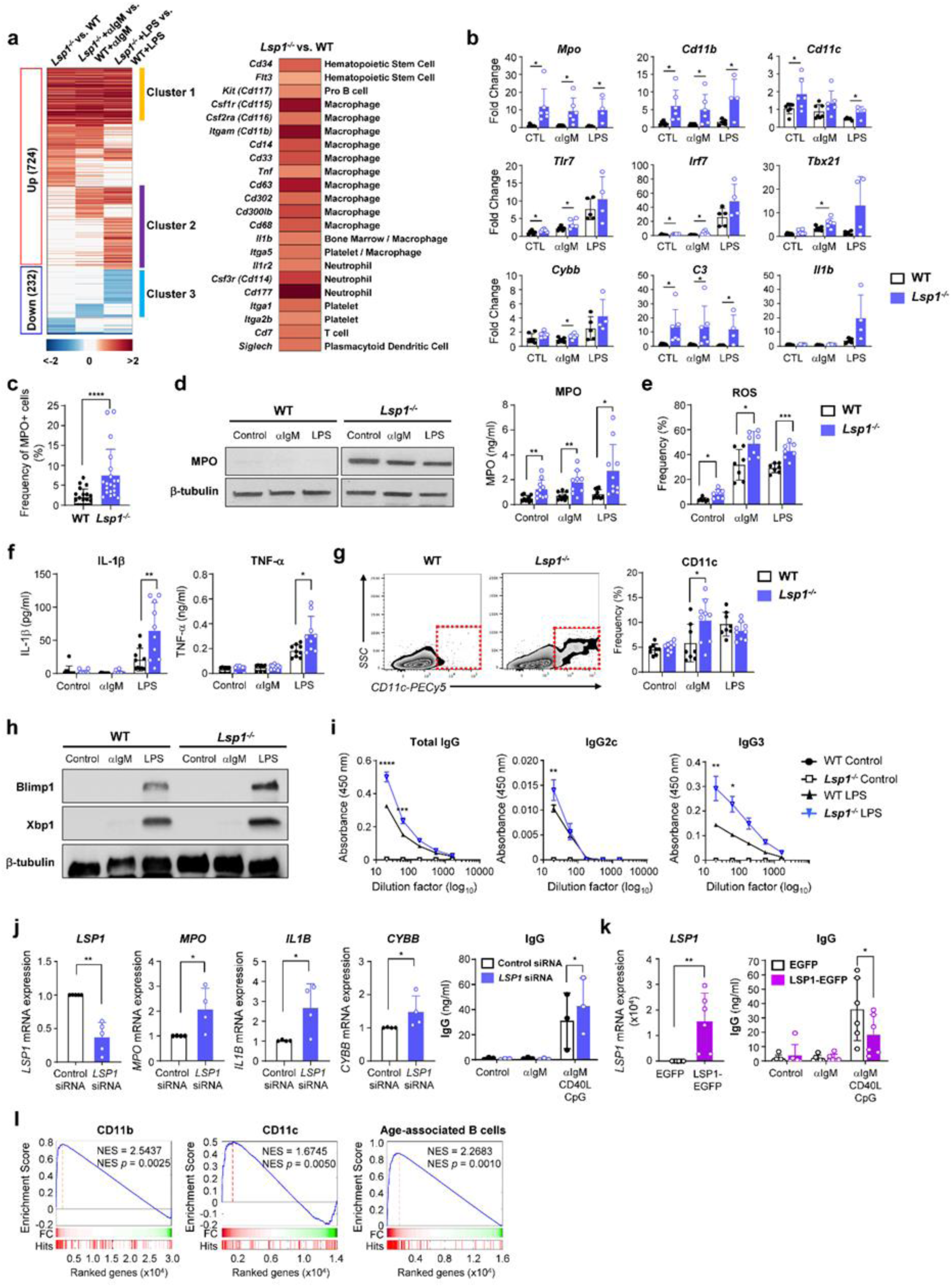
*Lsp1*-regulated B cells (LRBs) with both myeloid gene expression profiles and high Ab-producing ability. **a,** Heatmap displaying the DEGs between WT and *Lsp1^-/-^* B cells stimulated with αIgM or LPS for 4 hours. Red and blue indicate up- and downregulation, respectively (left panel). The right panel shows the hematopoietic cell markers upregulated by *Lsp1* deficiency. **b**, Upregulation of myeloid gene expression in *Lsp1^-/-^* B cells, as determined by qRT‒PCR; the fold changes were calculated using the 2^-ΔΔCt^ method. **c,** Percentage of MPO^+^ cells in splenic B cells freshly isolated from WT and *Lsp1^-/-^* mice (n=20 per group), as determined by immunofluorescence staining with anti-B220 (green) and anti-MPO (red) Abs. **d**-**i,** Splenic B cells isolated from WT and *Lsp1^-/-^* mice were stimulated with αIgM or LPS for 24 hours (**d-f**), 7 days (**g**), or 5 days (**h,i**). **d**, Immunoblot (left panel) and ELISA (right panel, n=10 per group) analyses of MPO expression. **e**, Intracellular ROS production (n=7 per group) by flow cytometry using H_2_DCFDA. **f**, Production of IL-1β and TNF-α (n=9∼10) by ELISA. **g**, Percentage of CD11c^+^ cells in WT and *Lsp1^-/-^* B cells (n=9∼10) by flow cytometry. A representative zebra plot for CD11c^+^ cells (red dotted box) is presented on the left. **h**, Immunoblotting for Blimp1 and Xbp1, which are representative TFs involved in B-cell differentiation. **i**, ELISA for IgG, IgG2c, and IgG3 produced by B cells stimulated with LPS (n=3 per group). **j,k,** *LSP1*-regulated gene expression and IgG production in human B cells. Sorted human B cells were electroporated with *LSP1* siRNA (**j**) or an LSP1-EGFP plasmid (**k**). After 24 hours, the mRNA expression levels of *LSP1*, *MPO*, *IL1B*, and *CYBB* in the cells were determined via qRT‒PCR. IgG production by B cells stimulated with αIgM, CD40L, and CpG for 5 days was determined via ELISA. **l,** GSEA plots demonstrating significant enrichment between the *Lsp1*-regulated genes in Clusters 1 and 2 and the genes (up- and downregulated) defining CD11b^+^ B cells, CD11c^+^ B cells, or ABCs. NES=normalized enrichment score. The data in the bar graphs are presented as the means ± SDs of at least two independent experiments. *P* values were determined by multiple unpaired two-tailed *t* tests (**b** to **g**), two-way ANOVA with Sidak’s multiple comparisons (**i**, right panels in **j** and **k**), and the Mann‒Whitney *U* or unpaired *t* test (left panels in **j** and **k**). * *P* < 0.05; ** *P* < 0.01; *** *P* < 0.001; and **** *P* < 0.0001.

### Lsp1-regulated B cells (LRBs) exhibit chimeric functions defined by both myeloid and lymphoid characteristics

Based on the transcriptome profiling results, we sought to determine whether *Lsp1* deficiency functionally bestows the myeloid phenotype on B lymphocytes. In contrast to lymphocytes, myeloid cells are able to produce oxidative enzymes such as myeloperoxidase (MPO) and reactive oxygen species (ROS)^18^. In particular, MPO is primarily released by activated macrophages and neutrophils but is rarely produced by B cells and functions as a tissue damage molecule to protect the host against pathogens^17^. Moreover, since MPO is one of the most highly upregulated myeloid genes in *Lsp1*^-/-^ B lymphocytes (**Extended Data Fig. 5b**), we first tried to validate elevated MPO expression in *Lsp1*^-/-^ B cells. As shown in **Fig. 2c**, the percentage of MPO-expressing B220^+^ B cells was higher in freshly isolated *Lsp1*^-/-^ B cells than in WT (*Lsp1*^-+/+^) B cells (**Fig. 2c** and **Extended Data Fig. 6a**). After stimulation with αIgM or LPS, *Lsp1*^-/-^ B lymphocytes also expressed and secreted increased amounts of MPO (**Fig. 2d**). Next, ROS production in the absence or presence of αIgM and LPS was determined, and the ROS production was also found to be higher in *Lsp1*^-/-^ B lymphocytes than in WT B lymphocytes (**Fig. 2e**), which was consistent with the increase in the expression levels of *Cybb*, the prototypical NADPH oxidase ^19^, in *Lsp1*^-/-^ B cells (**Fig. 2b**). Another characteristic of myeloid cells is the production of proinflammatory cytokines at high levels and the expression of the cell surface marker CD11c. As expected, *Lsp1*^-/-^ B cells produced more IL-1β and TNFα and exhibited higher CD11c expression on their surface when stimulated with LPS and αIgM, respectively (**Fig. 2f** and **2** **g**). In summary, these data demonstrated that *Lsp1*-deficient B cells have a myeloid phenotype and can produce high levels of myeloperoxidase (MPO), reactive oxygen species (ROS), and proinflammatory cytokines so that they can accelerate tissue damage and kill pathogens.

We wondered whether myeloid-like *Lsp1-*deficient B cells still retained the ability to produce Abs as classic B cells. To address this issue, we first measured the expression of Blimp1 and Xbp1, key transcription factors that regulate the differentiation of WT and *Lsp1*^-/-^ B cells to Ab-producing plasma cells^20^. The results showed that the expression of Blimp and Xbp1 was higher in *Lsp1*^-/-^ B cells stimulated with LPS for 5 days (**Fig. 2h**). In parallel, the total IgG and IgG3 levels, but not the IgM level, were markedly higher in *Lsp1*^-/-^ B cells stimulated with LPS than in WT B cells under the same stimulatory conditions (**Fig. 2i** and **Extended Data Fig. 6b**). This increase in Ab production was not related to B-cell survival or proliferation (**Extended Data 7a** and **7b**). Taken together, these findings, together with those of earlier reports^20^, demonstrate that Ab production is not reduced but rather strengthened in *Lsp1*^-/-^ B cells, which seems to be related to the upregulated expression of Blimp1 and Xbp1.

LSP1 upregulation of myeloid gene expression was reproduced in human B cells. As shown in **Fig. 2j**, the knockdown of *LSP1* transcripts via siRNA for 24 hours substantially upregulated the expression of myeloid genes, including *MPO*, *IL1B*, and *CYBB,* in the peripheral B cells of healthy subjects. Moreover, after combined stimulation with αIgM, CD40L, and CpG, *LSP1* siRNA-transfected B cells produced a greater amount of total IgG than control siRNA-transfected B cells, which was similar to the data obtained for mouse B cells (**Fig. 2j**). Conversely, when the *LSP1* gene was overexpressed in human B cells using the LSP1-EGFP plasmid, total IgG production was markedly suppressed (**Fig. 2k**). The expression of *MPO* and *IL1B* mRNA was undetectable in *LSP1*-expressing B cells (data not shown). Taken together, our mouse and human data demonstrated that *LSP1-*deficient B cells exhibit increased myeloid gene expression and strengthened Ab responses. Therefore, we referred to this subset of ‘chimeric B cells’ possessing both myeloid and lymphoid characteristics as LRBs.

We next compared the gene expression profiles of LRBs with those of special B-cell subsets expressing the myeloid markers CD11b and/or CD11c, as previously reported^21,3,22^. To this end, we collected gene expression datasets from three independent studies describing the characteristics of 1) CD11b^+^ B cells^21^, 2) CD11c^+^ B cells^3^, and 3) ABCs^22^ using the GEO database (see **Methods**). Gene set enrichment analysis (GSEA) was subsequently conducted, revealing a significant correlation between the *Lsp1*-regulated genes in clusters 1 and 2 and genes defining CD11b^+^ B cells, CD11c^+^ B cells, or ABCs (**Fig. 2l**). These data suggest that LRBs exhibit considerable similarity in terms of gene expression patterns to those of the other three types of myeloid B cells, suggesting that LSP1 functions as a regulator for the generation of ABCs.

### LRBs exhibit myeloid morphology with strong phagocytic activity

Identification of the upregulated myeloid-specific genes and functions of LRBs raised the interesting question of whether LRBs actually have myeloid morphology and phagocytic activity. Strikingly, upon immunofluorescence staining with phalloidin, αIgM-activated *Lsp1*^-/-^ B cells underwent great morphological changes mimicking myeloid cells, including a rich cytoplasm and pseudopod formation, in sharp contrast with αIgM-activated WT B cells, which maintained a large round nucleus and a thin cytoplasmic membrane (**Fig. 3a**). Along with these findings, the level of phalloidin staining was also substantially increased in *Lsp1*^-/-^ B cells. Transmission electron microscopy (TEM) analysis revealed ultrastructural changes in activated *Lsp1*^-/-^ B cells. In the resting state, both *Lsp1*^-/-^ and WT B cells had similar morphologies, with small and round shapes and a varying number of small dendrites from the cells; a large proportion of inactive nuclear chromatin (termed heterochromatin); and few cytoplasmic organelles, which are all typical lymphocyte morphologies, as presented in **Fig. 3b**. However, upon activation of the BCR signaling pathway with αIgM, *Lsp1*^-/-^ B cells became dramatically irregular in shape and exhibited multiple pseudopods (**Fig. 3b** and **Extended Data Fig. 8a**), which was consistent with the immunostaining data. Moreover, the proportion of uncondensed nuclear chromatin (termed euchromatin) and the number of rough endoplasmic reticulum were markedly greater in *Lsp1*^-/-^ B cells stimulated with αIgM and LPS, respectively (**Fig. 3b** and **Extended Data Fig. 8b)**, which suggested that compared to WT B cells, *Lsp1*^-/-^ B cells are transcriptionally and translationally active. Interestingly, after WT B cells were stimulated with R848, which is known to induce the production of ABCs^22^, a round shape of the cells was converted to myeloid morphology, similar the morphology of *Lsp1*^-/-^ B cells after IgM stimulation (**Fig. 3c**). Therefore, we concluded that *Lsp1* deficiency causes lymphoid-to-myeloid morphological changes in B cells upon BCR activation.

**Fig. 3.**
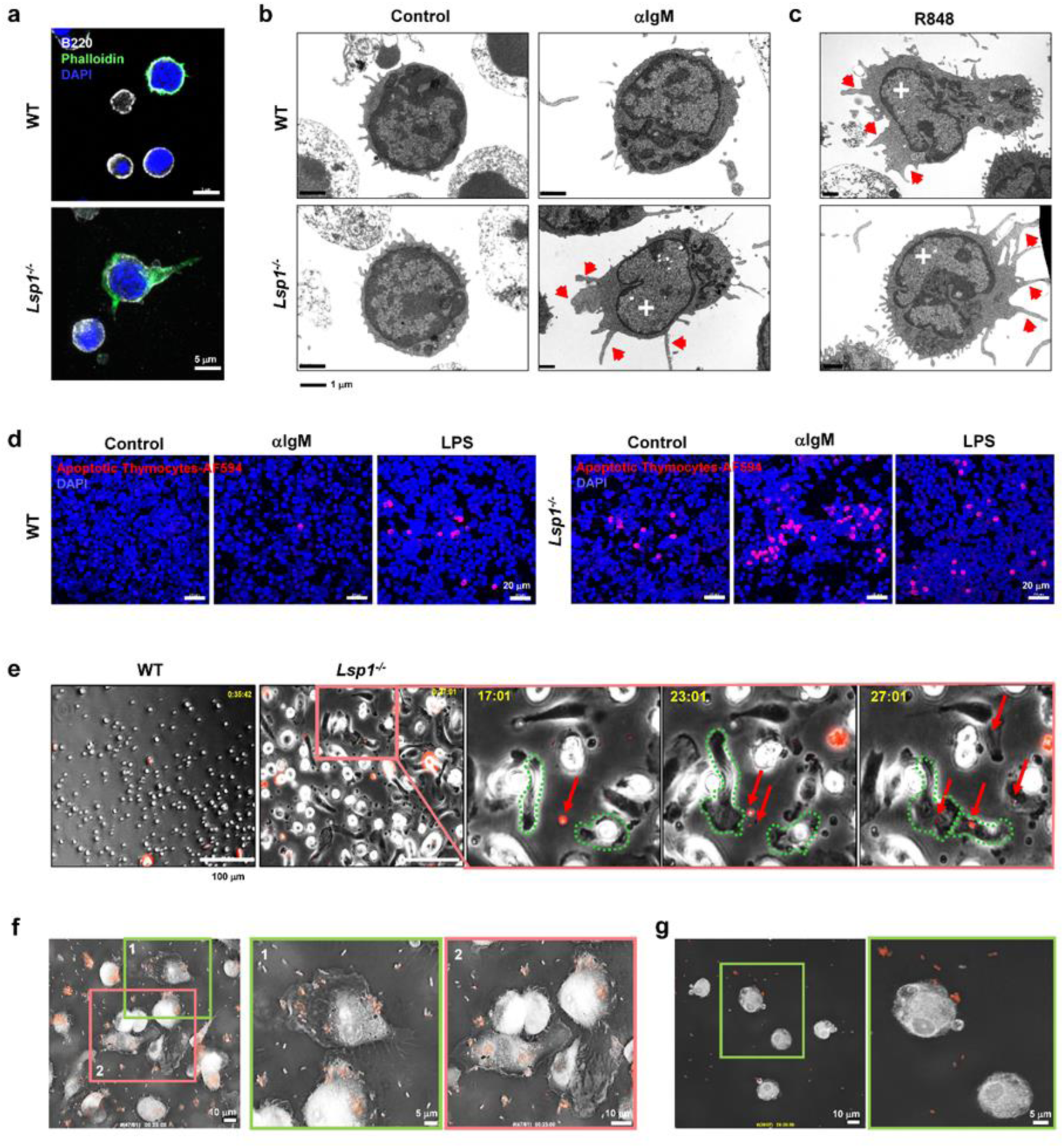
Myeloid morphology and phagocytic activity of B cells induced by *Lsp1* deficiency. **a,** Immunofluorescence staining of F-actin in B cells stimulated with αIgM-for 5 days using an anti-B220 Ab (white) and phalloidin (green). DAPI (blue) was used for nucleic acid staining. Scale bar, 5 μm. **b, c,** TEM images of B cells. Mouse splenic B cells stimulated with αIgM Ab (**b**) or R848 (**c**) for 3 days were subjected to ultrastructural analysis via TEM. The red arrows indicate pseudopodia, and the white crosses indicate euchromatin. Scale bar, 1 μm. **d,** Confocal images for detecting efferocytic activity. Mouse B cells were stimulated with αIgM or LPS for 11 days and treated with pHrodo-Red-labeled apoptotic thymocytes for 30 minutes. DAPI was used for nucleic acid staining. The pink color indicates the phagocytosed cells. Scale bar, 20 μm. The data are representative of more than three independent experiments. **e, f,** Phagocytic activity of αIgM-stimulated *Lsp1^-/-^* B cells. *Lsp1^-/-^* B cells were stimulated with αIgM for 11 days, after which the cells were treated with Alexa Fluor 594-labeled *E. coli* particles. *Lsp1^-/-^* B cells were continuously observed for 30 minutes for phagocytic activity and morphological changes via live-cell imaging with a LIONHeart LX (**e**) or TomoCube Holotomographic microscope (**f**). The red or green rectangular area in (**e**) and (**f**) is magnified on the right, which highlights the morphological changes that occur in *Lsp1^-/-^*B cells during the phagocytosis of *E. coli* particles (red arrows) in a time-dependent manner. In the magnified images in (**e**), the cells dotted in green were tracked over time**. g**, Inhibition of phagocytic activity by an actin polymerization inhibitor (cytochalasin D). After stimulating *Lsp1^-/-^* B cells with αIgM for 11 days, they were pretreated with 2.5 μM cytochalasin D for 30 minutes, after which Alexa Fluor 594-labeled *E. coli* particles (red dots) were added to the cells. Cellular movement was recorded for 30 minutes with a TomoCube Holotomographic microscope.

The myeloid morphology detected in activated *Lsp1*^-/-^ B cells prompted us to compare the phagocytic activity of *Lsp1*^-/-^ and WT B cells (**Fig. 3c**). When apoptotic thymocytes labeled with pHrodo-Alexa Fluor 594 were added to the two types of B cells for 30 minutes, the *Lsp1*^-/-^ B cells exhibited higher levels of fluorescence staining (pink) than the WT B cells, particularly under αIgM-stimulated conditions (**Fig. 3d**), suggesting an increase in the phagocytic activity of self-antigens in response to *Lsp1* deficiency. We also examined the ability of *Lsp1*^-/-^ B cells to phagocytose foreign pathogens. Surprisingly, immediately after the addition of *E. coli* particles, the majority of αIgM-stimulated *Lsp1*^-/-^ B cells had adhered to the culture plate and aggressively migrated toward *E. coli*, and some of these cells actively engulfed and internalized *E. coli* via elongated pseudopodia (**Fig. 3e** and **Extended Data Video 1**), which was exactly what we commonly observed in activated macrophages exposed to pathogens. In contrast, such morphologic changes and phagocytic activity were hardly detected in the αIgM-activated WT B cells cocultured with *E. coli*. (**Fig. 3e** and **Extended Data Video 2**).

To scrutinize the phagocytic process of *Lsp1*^-/-^ B cells in detail and to determine the involvement of actin polymerization in this process, we utilized the “Tomocube system”, which employs holotomographic techniques using a digital micromirror device. Under the same conditions as in **Fig. 3e**, we confirmed that after the addition of *E. coli* particles, the IgM-stimulated *Lsp1*^-/-^ B cells became larger and more irregular in shape, actively generated pseudopodia, and subsequently engulfed and internalized the *E. coli* particles (**Fig. 3f** and **Extended Data Video 3**). In sharp contrast, the addition of cytochalasin D, an actin polymerization inhibitor, completely blocked the morphological changes and active movement of αIgM-stimulated *Lsp1*^-/-^ B cells in response to *E. coli*, as indicated by the decreased phagocytic activity. Moreover, the cells did not phagocytose the *E. coli* particles (**Fig. 3g** and **Extended Data Video 4**), demonstrating that actin polymerization is required for the phagocytic ability of LRBs.

Taken together, these findings indicate that activated LRBs exhibit a distinct myeloid morphology with robust phagocytic activity against self-antigens and foreign antigens, suggesting that LSP1, as an actin remodeling regulator, plays a central role in the phagocytic activity and myeloid morphology of this B-cell subset.

### The chimeric function of LRBs is mediated by the PKCβ-CEBPβ pathway

We next questioned which signaling pathways are responsible for the dual myeloid and lymphoid functions of LRBs. To this end, we first carried out a master regulator analysis (MRA) using a public database to define the interactions between transcription factors (TFs) and their target genes (see **Methods**). The three sets of master regulators were predicted through TF enrichment analysis using the 369 upregulated DEGs in *Lsp1*^-/-^ B cells with or without αIgM or LPS treatment (**Fig. 4a**, left panel). Notably, CCAAT/enhancer binding protein (CEBP)α and CEBPβ were enriched in all three sets (**Fig. 4a**, left panel). In particular, when the number of target genes was checked, CEBPβ was found to interact with more than 50% of the DEGs (**Fig. 4a**, right panel), suggesting that CEBPβ predominantly governs the majority of the genes upregulated by *Lsp1* deficiency. By contrast, CEBPα interacted with only about 10% of the DEGs. This predominance of CEBPβ could also be observed in the molecular network analysis of the DEGs involved in hematopoietic cell lineage markers, phagocytosis, and cytokine/chemokine pathways (**Fig. 4b**). Taken together, these findings strongly suggested that CEBPβ is the key TF mediating the chimeric functions of LRBs.

**Fig. 4.**
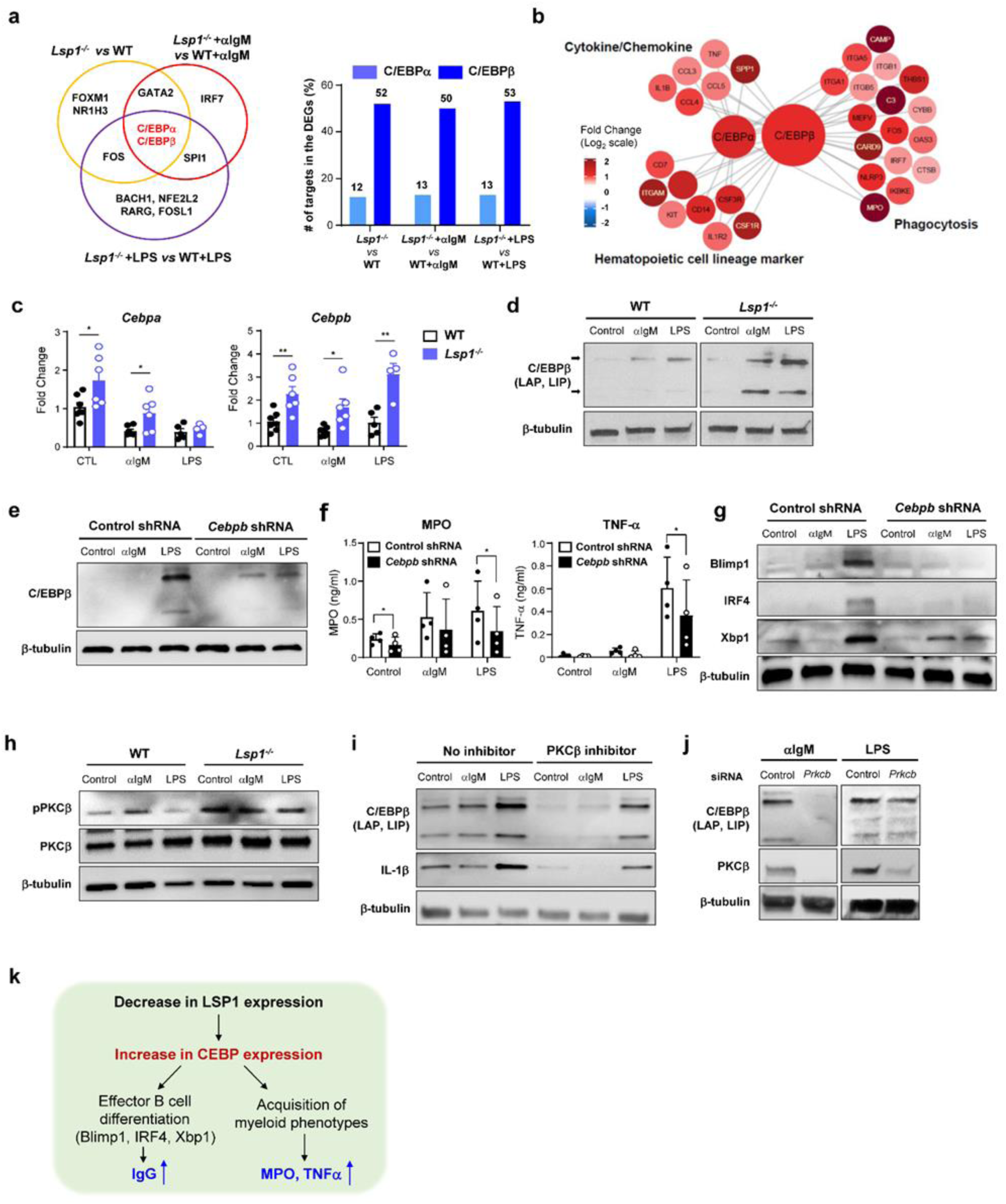
Requirement of the PKCβ-C/EBPβ pathway for the chimeric functions of LRBs. **a,** Venn diagram displaying transcription factors (TFs) that have a high potential to regulate the 369 upregulated DEGs in *Lsp1^-/-^* B cells with or without αIgM or LPS treatment (left panel). The bar graph on the right depicts the numbers of C/EBP target genes among the DEGs in *Lsp1^-/-^* B cells. **b,** PPI network representing the interactions between C/EBPα/β and the 31 C/EBP target genes involved hematopoietic cell lineage markers, cytokines/chemokines, and phagocytosis. The color of the nodes indicates the fold change in the genes affected by *Lsp1* deficiency. **c, d,** Quantitative RT‒PCR analysis of *Cebpa* and *Cebpb* mRNA expression (n=4∼7; **c**) and immunoblotting for C/EBPβ protein expression (**d**). *Lsp1^-/-^*versus WT B cells were stimulated with αIgM or LPS for 4 hours (**c**) or for 24 hours (**d**). **e, f,** *Lsp1^-/-^* B cells were transduced with control or *Cebpb* shRNAs, and the cells were then stimulated with αIgM or LPS for 24 hours for C/EBPβ immunoblotting (**e**) and MPO and TNF-α ELISAs (n=4; **f**). **g,** Control or *Cebpb* shRNA-treated *Lsp1^-/-^* B cells were stimulated with αIgM or LPS for 5 days, after which Blimp1, IRF4, and Xbp1 expression was determined via immunoblot assay. **h,** Immunoblots of phosphorylated PKCβ (pPKCβ) in *Lsp1^-/-^* versus WT B cells. Splenic B cells were stimulated with αIgM or LPS for 5 minutes. **i, j,** Effects of PKCβ inhibition on C/EBPβ and IL-1β expression in *Lsp1^-/-^* B cells stimulated with αIgM or LPS. (**i**) *Lsp1^-/-^* B cells were pretreated with the PKCβ inhibitor ruboxistaurin (4 μM) for 1 hour and stimulated with αIgM or LPS for 24 hours. (**j)** *Lsp1^-/-^* B cells were stimulated with αIgM or LPS for 24 hours and then electroporated with control or *Prkcb* siRNA for an additional 24 hours. The expression levels of C/EBPβ, its target gene IL-1β, and PKCβ were determined via an immunoblot assay. **k,** Proposed model for the dual regulation of B-cell functions by LSP1. The data in the bar graphs are presented as the means ± SDs of at least three independent experiments. *P* values were determined by multiple unpaired *t* tests (**c**) and paired *t* tests (**f**). * *P* < 0.05; ** *P* < 0.01.

In support of this notion, *Lsp1*^-/-^ B cells presented higher levels of CEBPβ mRNA and protein expression than WT B cells after αIgM or LPS stimulation (**Fig. 4c** and **4d**). Moreover, the shRNA-mediated knockdown of CEBPβ transcripts resulted in a substantial decrease in the secretion of MPO and TNF-α, the key myeloid genes from *Lsp1^-/-^* B cells stimulated with LPS (**Fig. 4e** and **4f**). Regarding Ab-producing ability, the expression of Blimp1, IRF4, and XBP1, the essential TFs for differentiation into Ab-producing cells^20^, was also markedly downregulated in CEBPβ shRNA-treated *Lsp1*^-/-^ B cells (**Fig. 4g**). Taken together, the results of the master regulator analysis, network analysis, and knockdown experiments revealed that CEBPβ appears to be indispensable for LRBs to acquire both myeloid cell functions and elevated Ab-producing abilities.

However, how LSP1 deficiency is related to CEBP upregulation has not been determined. Previous studies have reported that LSP1 represses the activation of PKCβ in B cells^11^ and that PKC activation induces CEBPβ expression in promyelocytic cells^23^, which raises the possibility of a LSP1-PKCβ-CEBPβ axis in LRBs. As expected, the level of phosphorylated PKCβ (pPKCβ) was higher in *Lsp1*^-/-^ B cells than in WT B cells (**Fig. 4h**). Moreover, pretreatment of *Lsp1*^-/-^ B cells with the PKCβ inhibitor ruboxistaurin substantially downregulated CEBPβ expression as well as IL-1β expression (**Fig. 4i**). Moreover, the ERK inhibitor PD98059 did not affect CEBPβ expression (**Extended Data Fig. 9**). In addition, the transfection of *Lsp1*^-/-^ B cells with PKCβ siRNAs dramatically downregulated CEBPβ expression in the presence of αIgM or LPS (**Fig. 4j**). These results, together with earlier reports^11,23^, suggest that LSP1 regulates CEBPβ expression by acting on PKCβ.

In summary, LSP1 deficiency upregulates CEBPβ expression via the activation of PKCβ, which may be necessary for the chimeric function of LRBs (**Fig. 4k**).

### Lsp1 deficiency exacerbates pristane-induced lupus in mice

Given the bioinformatics and *in vitro* data, we sought to validate our hypothesis *in vivo*. To this end, we established a mouse model of pristane-induced lupus, which is a representative B-cell-dependent autoimmune disease^24^ characterized by pulmonary vasculitis culminating in diffuse lung hemorrhage, glomerulonephritis, and autoantibody generation^25^. Two weeks after pristane injection, LSP1 expression was partially downregulated, whereas CEBPβ and MPO expression was considerably upregulated in splenic B cells (**Fig. 5a**). Interestingly, the expression of ETS1, an upstream TF of *Lsp1*^26^, was also partially downregulated (**Fig. 5a**), which suggests the possible involvement of ETS1 in regulating LSP1 in pristane-induced lupus. Additionally, the percentages of CD11b^+^CD11c^+^ (double-positive) B cells and CD19^-^CD138^+^ cells were significantly increased (**Fig. 5b**), indicating that pristane induced the expansion of ABCs and Ab-producing plasma cells, respectively. Taken together, these findings suggest that pristane-induced autoimmunity may lead to increases in CEBPβ expression levels, MPO release, and ABC production along with a decrease in LSP1 expression levels.

**Fig. 5.**
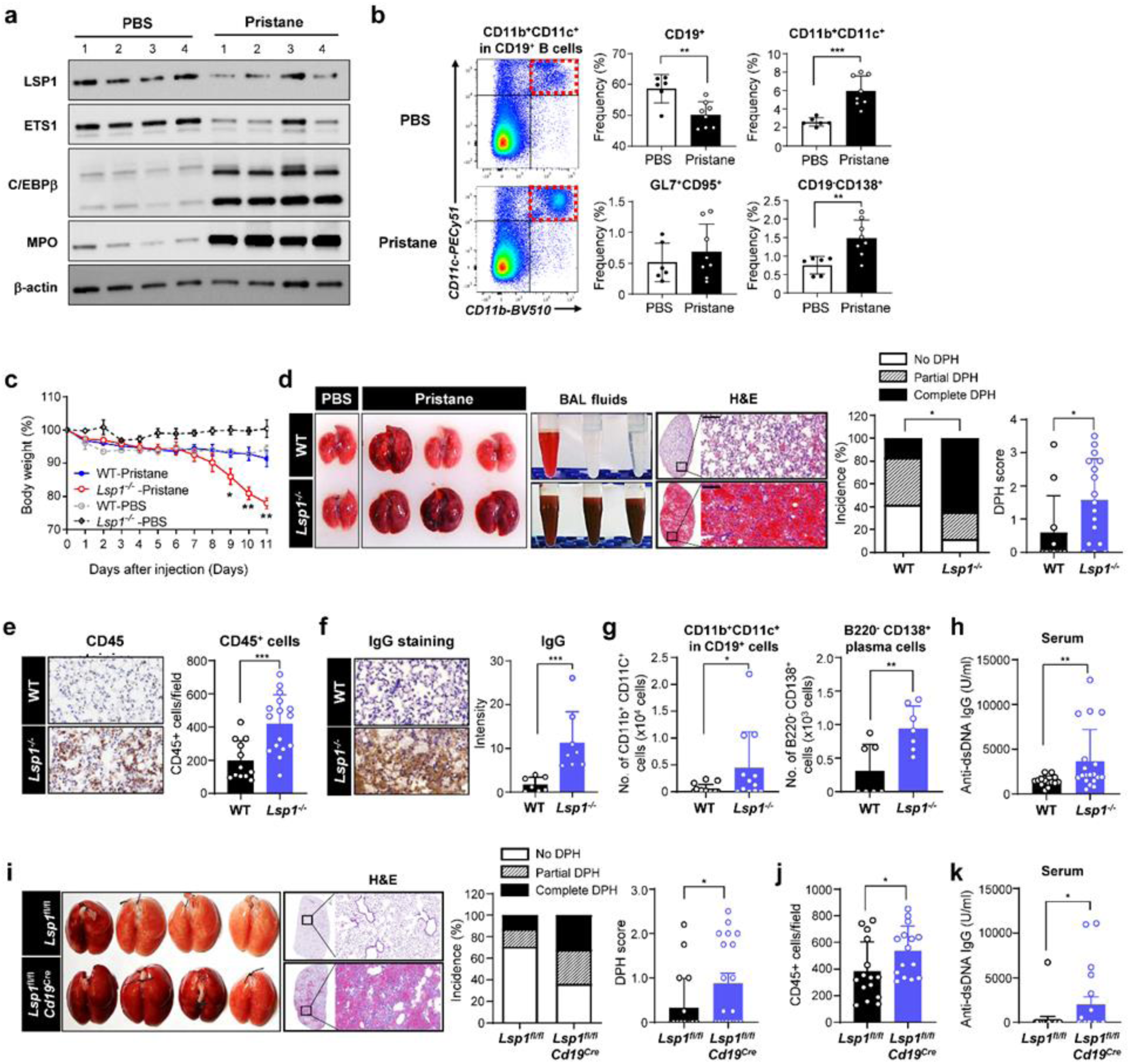
Deterioration of pristane-induced lupus by *Lsp1* deficiency. **a,b,** Effects of pristane on the expression of LSP1 and its target genes. WT mice were intraperitoneally injected with 500 μl of pristane or PBS as a control (n=4 per group). Two weeks later, the expression of LSP1, its upstream regulator ETS1, and its downstream target genes C/EBPβ and MPO in splenic B cells was determined. (**a**) Immunoblot analysis. The numbers indicate the individual mice. (**b**) Flow cytometry. The percentages of B-cell subpopulations, including total CD19^+^ B cells, GC B cells (GL7^+^CD95^+^ in CD19^+^ cells), ABCs (CD11b^+^CD11c^+^ in CD19^+^ cells), and plasma cells (CD19^-^CD138^+^ cells), were analyzed. A representative pseudocolor plot (left panel) shows the ABC population in the rectangles with red dots in the PBS (n=6) and pristane (n=8) groups. **c-h,** Aggravation of pristane-induced lupus by *Lsp1* deficiency. WT (n=7) and *Lsp1^-/-^* (n=8) mice were intraperitoneally injected with pristane. (**c**) Changes in body weight. (**d**) Diffuse pulmonary hemorrhage (DPH) severity in the lungs and bronchoalveolar lavage fluids (BALFs) of WT (n=12) and *Lsp1^-/-^* (n=17) mice was evaluated via gross inspection. DPH incidence and DPH scores of the two groups were also determined by H&E staining. Infiltration of CD45^+^ leukocytes (**e**) and IgG expression (**f**) in the lungs of WT and *Lsp1^-/-^*mice were assessed by immunohistochemistry using anti-CD45 and anti-IgG Abs. The numbers of CD45^+^ cells were manually counted from 5 fields per slide. The intensity of IgG expression in 3 fields per lung tissue sample was analyzed using ImageJ. **g**, The number of ABCs and plasma cells in the BALF of WT and *Lsp1^-/-^* mice was assessed via flow cytometry. **h**, ELISA for anti-dsDNA IgG levels in the sera of WT (n=14) and *Lsp1^-/-^*(n=19) mice. **i-k,** Acceleration of pristane-induced lupus by B-cell-specific depletion of *Lsp1*. *Lsp1^fl/fl^* and *Lsp1^fl/fl^ Cd19^cre/-^*mice were intraperitoneally injected with pristane. (**i**) DPH severity in the lungs and BALF of the two groups of mice (n=19 per group). (**j)** Infiltration of CD45^+^ cells in the lungs of the two groups (n=15 per group). (**k**) Anti-dsDNA IgG production in the sera of *Lsp1^fl/fl^* (n=20) and *Lsp1^fl/fl^ Cd19^cre/-^*(n=19) mice. The data in the bar graphs are presented as the means ± SDs of at least three independent experiments. *P* values were determined by the Mann‒Whitney *U* test (**b, d** to **k,** except for incidence data), two-way ANOVA with Sidak’s multiple comparisons (**c**), and Fisher’s exact test (percentage of incidence in **d** and **i**). * *P* < 0.05; ** *P* < 0.01; *** *P* < 0.001.

To further probe the pathological relevance of LSP1, we injected pristane into *Lsp1*^-/-^ and WT mice. As shown in **Fig. 5c**, *Lsp1*^-/-^ mice exhibited a more dramatic decrease in body weight from 8 to 11 days after pristane injection than WT mice (**Fig. 5c**). Strikingly, on day 11, the bronchoalveolar lavage fluid (BALF) and gross morphology of the lung were completely hemorrhagic (100%) in *Lsp1*^-/-^ mice and only partially hemorrhagic (14.3%) in WT mice (**Fig. 5d**). Consistently, the diffuse pulmonary hemorrhage (DPH) score, which was determined by assessing the histopathology of the lung, was significantly greater in *Lsp1*^-/-^ mice than in WT mice (**Fig. 5d**). Moreover, the number of CD45^+^ (a marker of leukocytes) cells and the IgG positivity per field in the lung tissues were higher in *Lsp1*^-/-^ mice than in WT mice (**Fig. 5e** and **5f**), indicating that the stronger pulmonary inflammation and Ab production in association with *Lsp1* deficiency were elicited by pristane injection. Importantly, the percentages of CD11b^+^CD11c^+^ CD19^+^ B cells (ABCs) and B220^-^CD138^+^ plasma cells were higher in the BALF of *Lsp1*^-/-^ mice than in that of WT mice (**Fig. 5g**), which suggested that *Lsp1* deficiency resulted in both a myeloid phenotype and greater Ab production of B cells (LRB generation) at the injury site in *Lsp1*^-/-^ mice. Moreover, the number of CD11b^+^ cells and CD11b^+^ Ly6C^+^ monocytes was also greater in the BALF of *Lsp1*^-/-^ mice (**Extended Data Fig. 10a** and **10b**). In parallel, the levels of anti-dsDNA Abs, which are a hallmark of lupus pathology^25^, were also elevated in the sera of pristane-induced *Lsp1*^-/-^ mice (**Fig. 5h**).

To clarify that B cells were specifically involved in the observed role of LSP1 in pristane-induced lupus, we generated conditional knockout (KO, haploinsufficient) mice lacking *Lsp1* specifically in B cells using the Cre/loxP system. As expected, in *Lsp1^fl/fl^Cd19^Cre/-^*mice, *Lsp1* mRNA and protein levels were reduced only in B cells and not in other cell types, such as T lymphocytes and macrophages, confirming the specific depletion of *Lsp1* in B cells (**Extended Data Fig. 11**). When *Lsp1^fl/fl^Cd19^Cre/-^*mice were subjected to pristane-induced lupus, they exhibited more severe pulmonary hemorrhages and greater infiltration of CD45^+^ leukocytes in the lungs than *Lsp1^fl/fl^* mice (**Fig. 5i** and **5j**), demonstrating the decisive effect of LSP1 in B cells on pristane-induced lupus. Consistent with these findings, anti-dsDNA Ab levels were significantly higher in the sera of pristane-induced *Lsp1^fl/fl^Cd19^Cre/-^*mice than in those of control littermates (**Fig. 5k**). Taken together, the results of the B-cell-specific *Lsp1* KO mice experiments confirmed that LRBs accelerate pristane-induced lupus, promoting pulmonary vasculitis (hemorrhage) and autoantibody production.

### Downregulated LSP1 expression in lupus patients correlates with MPO expression in B cells, disease activity, and the IFN signature

Most studies examining the role of LSP1 in disease have been conducted in *Lsp1*-deficient mice, with few studies in human systems. To ascertain the clinical relevance of LRBs in human autoimmune diseases, we first measured LSP1 expression in B cells from patients with SLE, a classic autoimmune disease in which B cells, specifically ABCs, play a central role^27^. As shown in **Fig. 6a**, LSP1 expression was substantially downregulated in the peripheral B cells of patients with SLE compared to healthy donors, and it was negatively correlated with the systemic lupus erythematosus disease activity index (SLEDAI)^28^, which is the standard method for measuring SLE activity, as determined immediately after the isolation of peripheral blood. Moreover, the expression levels of LSP1 was also markedly lower in SLE B cells stimulated with αIgM, CD40L, and CpG (**Fig. 6b**) than in healthy B cells under the same conditions.

**Fig. 6.**
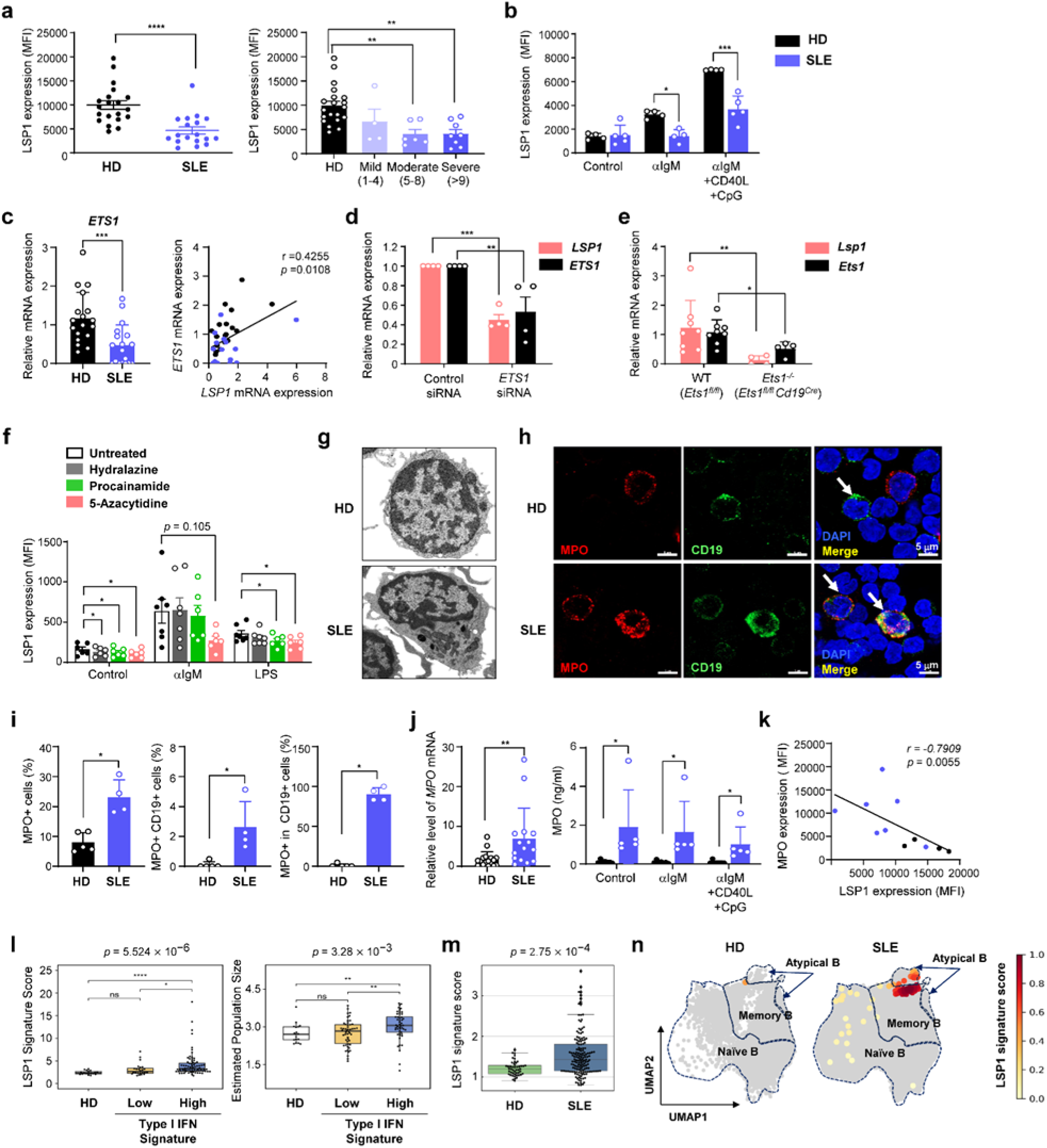
LSP1 expression in B cells and its correlation with ETS1 and MPO expression and disease activity in SLE patients. **a,** Flow cytometry was used to determine LSP1 expression levels in the peripheral B cells of healthy donors (HD, n=20) and SLE patients (n=18): MFI=mean fluorescence intensity. Disease activity was evaluated by the SLEDAI. **b,** LSP1 expression in cultured B cells from HDs (n=4) versus SLE B cells (n=4∼5) determined by flow cytometry. B cells were stimulated with αIgM, CD40L, and CpG for 3 days. **c,** qRT‒PCR analysis of *ETS1* mRNA expression in B cells from HDs (n=18) and SLE patients (n=17, left panel) and its correlation with *LSP1* mRNA expression in HDs (black circle) and SLE patients (blue circle, right panel). **d,e,** Regulation of LSP1 expression by ETS1. **d**, *ETS1* and *LSP1* mRNA expression in human B cells transfected with *ETS1* siRNA, as evaluated by qRT‒PCR (n=4). **e**, qRT‒PCR analysis of *Ets1* and *Lsp1* mRNA in B cells from WT (*Ets1^fl/fl^*, n=8) and *Ets1^-/-^* (*Ets1^fl/fl^C d19^Cre/-^*, n=4) mice. **f,** LSP1 expression in WT B cells (n=6∼7) stimulated with αIgM and LPS for 5 days in the presence or absence of (pretreated) 20 μM hydralazine, 20 μM procainamide, or 1 μM 5-azacytidine, as examined by flow cytometry. **g,** TEM images of B cells from HDs and SLE patients. **h, i,** Immunofluorescence staining of MPO in HDs and SLE B cells using anti-MPO (red) and anti-CD19 (green) Abs and DAPI (blue). **h**, Representative images of MPO^+^ B cells are indicated by white arrows. **i**, Percentage of MPO^+^ B cells determined by analyzing 6 fields per sample (n=5 for HDs, n=4 for SLE patients). **j,** qRT‒PCR and ELISA results for MPO. Left: *MPO* mRNA expression in HDs (n=17) versus SLE B cells (n=15). Right: ELISA for MPO production by B cells from HDs (n=7) and SLE patients (n=5) activated with the indicated stimuli for 3 days. **k,** Flow cytometry analysis of LSP1 and MPO expression in CD19^+^ B cells from HDs (black circle, n=4) and SLE patients (blue circle, n=7). **l,** Left: LSP1 signature in SLE patients and its correlation with the type I IFN signature: n=24 for the low IFN signature, n=75 for the high IFN signature, and n=181 for HD. Bulk RNA-seq data from SLE PBMCs were obtained from a public database (GSE72509). Right: Estimated population size of monocytic cells by cellular deconvolution analysis. Each dot represents an SLE patient or HD. **m**, LSP1 signature score in B cells from SLE patients versus HDs determined by pseudobulk RNA-seq analysis (GSE174188; s*ee* **Methods**). **n,** Uniform Manifold Approximation and Projection (UMAP) plot visualizing the B-cell subsets with high LSP1 scores (>0.8) in SLE patients. The data in the bar graphs are presented as the means ± SDs of at least two independent experiments. *P* values were determined by the Mann‒Whitney *U* test (left panels in **a**, **c**, and **j**; all data in **i**); one-way ANOVA with Dunnett’s multiple comparisons (right panel in **a**); two-way ANOVA with Sidak’s, Dunnett’s multiple comparison or multiple *t* test (**b**, **d**-**f**; right panel in **j**); and the Spearman correlation test (right panels in **c** and **k**). * *P* < 0.05, ** *P* < 0.01, *** *P* < 0.001, and **** *P* < 0.0001.

We next investigated which mechanisms drive the downregulation of LSP1 in lupus patients. Multiple genetic, epigenetic, and environmental factors have been associated with the pathogenesis of SLE^29^, and they might be causative factors for the downregulated LSP1 expression in SLE patients. To test this possibility, we first examined the relationship between LSP1 and ETS1 gene expression since ETS1 not only is a well-established genetic risk factor for SLE but also functions as an upstream regulator of *LSP1* transcription^26,30^. As previously reported^30^, *ETS1* mRNA expression in B cells was significantly lower in SLE patients than in healthy donors (**Fig. 6c**, left panel). Interestingly, *ETS1* expression was closely correlated with *LSP1* expression (**Fig. 6c**, right panel), which supports the findings of an earlier report^26^. Moreover, the transfection of *ETS1* siRNAs into human peripheral B cells significantly downregulated *LSP1* mRNA expression; in contrast, *LSP1* siRNAs only modestly downregulated *ETS1* mRNA expression in B cells (**Fig. 6d** and **Extended Data Fig. 12**). In parallel, *Lsp1* mRNA expression levels were markedly reduced in the B cells of the *Ets1*^-/-^ mice compared to those of their WT littermates (**Fig. 6e**). These results, along with those of previous reports^26^, show that the downregulated LSP1 expression in SLE patients may be associated with genetic defects in the *ETS1* gene.

A variety of environmental factors can trigger SLE, including drugs and viral infection that activate Toll-like receptor 7 (TLR7)^31^. To identify the environmental factors leading to LSP1 downregulation, we first treated mouse B cells with lupus-inducing drugs^32,33^, such as procainamide and hydralazine. Intriguingly, the two drugs significantly downregulated LSP1 expression in the absence of αIgM or LPS (**Fig. 6f**). Since hydralazine and procainamide are known to be competitive inhibitors of DNA methyltransferase^33^, another DNA methylation inhibitor, 5-azacytidine^34^, was also tested and found to strongly downregulate basal and αIgM- or LPS-stimulated LSP1 expression in B cells (**Fig. 6f**). Additionally, R848 injection into mice to stimulate TLR7 time-dependently downregulated LSP1 expression in B cells over 2 weeks, accompanied by an increase in CEBPβ and MPO expression levels. Similar to that observed in pristane-treated mice, the percentages of CD11b^+^ CD11c^+^ and CSF1R^+^ CD19^+^ cells were greater in the spleens of *Lsp1*^-/-^ mice after R848 injection (**Extended Data Fig. 13**). Collectively, these data suggest that lupus-inducing drugs and TLR7 stimulants can be additional causative factors of the downregulated LSP1 expression in patients with SLE.

We next questioned whether SLE B cells have the same morphological phenotype and molecular functions as LRBs. As shown in **Fig. 6g** and **Extended Data Fig. 14**, similar to those of LRBs, a considerable number of freshly isolated SLE B cells exhibited a markedly irregular shape with eccentric nuclei, profuse cytoplasm, and multiple pseudopods according to the TEM analysis; these features were rarely noted in healthy B cells. Moreover, the expression of MPO, the myeloid gene most highly upregulated in LRBs in association with phagosomes/phagocytosis (**Extended Data Fig. 5b**), was markedly upregulated in drug-naïve SLE B cells (**Fig. 6i**, left panel of **6j**), as determined by immunostaining and qRT‒PCR. Indeed, in contrast with healthy B cells (< 1% of MPO^+^ B cells), more than 90% of drug-naïve SLE B cells expressed MPO (right panel of **6i)**. Consistently, MPO production by B cells cultured in the presence of αIgM, CD40L, and/or CpG was also greater in SLE patients than in healthy controls (right panel of **6j**). A negative correlation was found between LSP1 and MPO expression (**Fig. 6k**), which is in line with the LRB data. Together, these findings revealed that the morphology of SLE B cells resembled that of LRBs, as they had a high capacity for producing MPO. Notably, oral administration of 20 μM of the MPO inhibitor PF06281355 every other day for 2 weeks significantly improved pulmonary hemorrhage in the lungs of *Lsp1^-/-^* mice with pristane-induced lupus (**Extended Data Fig. 15**). These results, together with the upregulated MPO expression in LRBs and SLE B cells, support the idea that MPO is one of the key effector molecules mediating lung pathology in lupus.

To further characterize the pathogenic relevance of LSP1 in SLE B cells, we selected 33 genes, namely, CEBPα, CEBPβ, and 31 target genes of CEBP, in *Lsp1*-deficient B cells (**Fig. 4b**) and designated them the **‘**LSP1 signature’. The LSP1 signature was scored based on the expression levels of the 33 genes and subsequently compared with the gene expression profile of SLE B cells, which was extracted from two public datasets^35,36^. As a result, the LSP1 signature score was much greater in SLE patients than in healthy controls, particularly in patients with a high type I interferon (IFN) signature (the left panel of **Fig. 6l** and **6** **m**), which is regarded as a therapeutic target and an indicator of disease activity in SLE patients^37^. Moreover, the LSP1 signature score correlated with the estimated population size of myeloid cells among peripheral mononuclear cells (right panel of **Fig. 6l**). Additionally, analysis of the single-cell RNA sequencing data revealed that the LSP1 signature was elevated in B cells of SLE patients compared to those of healthy donors (HDs) (**Fig. 6m**), and it was more prevalent in the atypical B cells of SLE patients^36^, akin to ABCs, than in naïve and memory B cells (**Fig. 6n**).

Overall, LSP1 downregulation can be induced by an *ETS1* gene defect, lupus-inducing drugs, and TLR7 ligands and is inversely correlated with MPO expression in B cells, disease activity, and the IFN signature in SLE patients.

### Retinoic acid ameliorates murine and human lupus through regulation of the LSP1-CEBPβ-MPO axis

The *in vitro*, *in vivo*, human pathology, and transcriptome profiling results indicate that LSP1 may be an attractive target for SLE treatment. Therefore, we hypothesized that a new drug designed to upregulate LSP1 expression could slow the progression of autoimmune diseases by reducing CEBPβ and MPO expression levels in LRBs. To test this possibility, we used a chemical-gene interaction database (CtdBase) and searched for chemicals that regulate LSP1 expression. Among the 106 chemicals reported to upregulate LSP1 expression, tretinoin, also known as all-trans retinoic acid (ATRA), was selected according to the criteria of the number of gene sets reported (counts), safety, and toxicity (**Extended Data Fig. 16**).

After splenic B cells from mice were treated with ATRA, LSP1 expression levels were found to increase in a dose-dependent manner (**Fig. 7a**, left panel). LSP1 protein expression, stimulated with αIgM or αIgM+CD40L, was further increased by ATRA (**Fig. 7a**, right panel). The mRNA expression levels of *Lsp1* as well as *Ets1* were also upregulated by ATRA, suggesting transcriptional regulation of both LSP1 and ETS1 by ATRA (**Fig. 7b**). With these data, we further examined the therapeutic effect of ATRA on pristane-induced lupus in mice. As shown in **Fig. 7c** and **7d**, intraperitoneal injection of 0.5 mg/kg ATRA (3 times a week) markedly alleviated lung hemorrhage and CD45^+^ cell infiltration in the lungs 2 weeks after pristane injection. Concurrently, ATRA injection robustly upregulated LSP1 expression while downregulating CEBPβ and MPO expression in the splenic B cells of mice with pristane-induced lupus (**Fig. 7e**). Moreover, the percentage of CD11b^+^CD11c^+^ CD19^+^ B cells in the spleen, but not that of CD19^+^ B cells, was decreased by ATRA injection in the same mouse model (**Fig. 7f**). Taken together, these *in vitro* and *in vivo* data indicate that ATRA reduces the generation of ABCs as well as the expression levels of CEBPβ and MPO by increasing LSP1 levels in B cells.

**Fig. 7.**
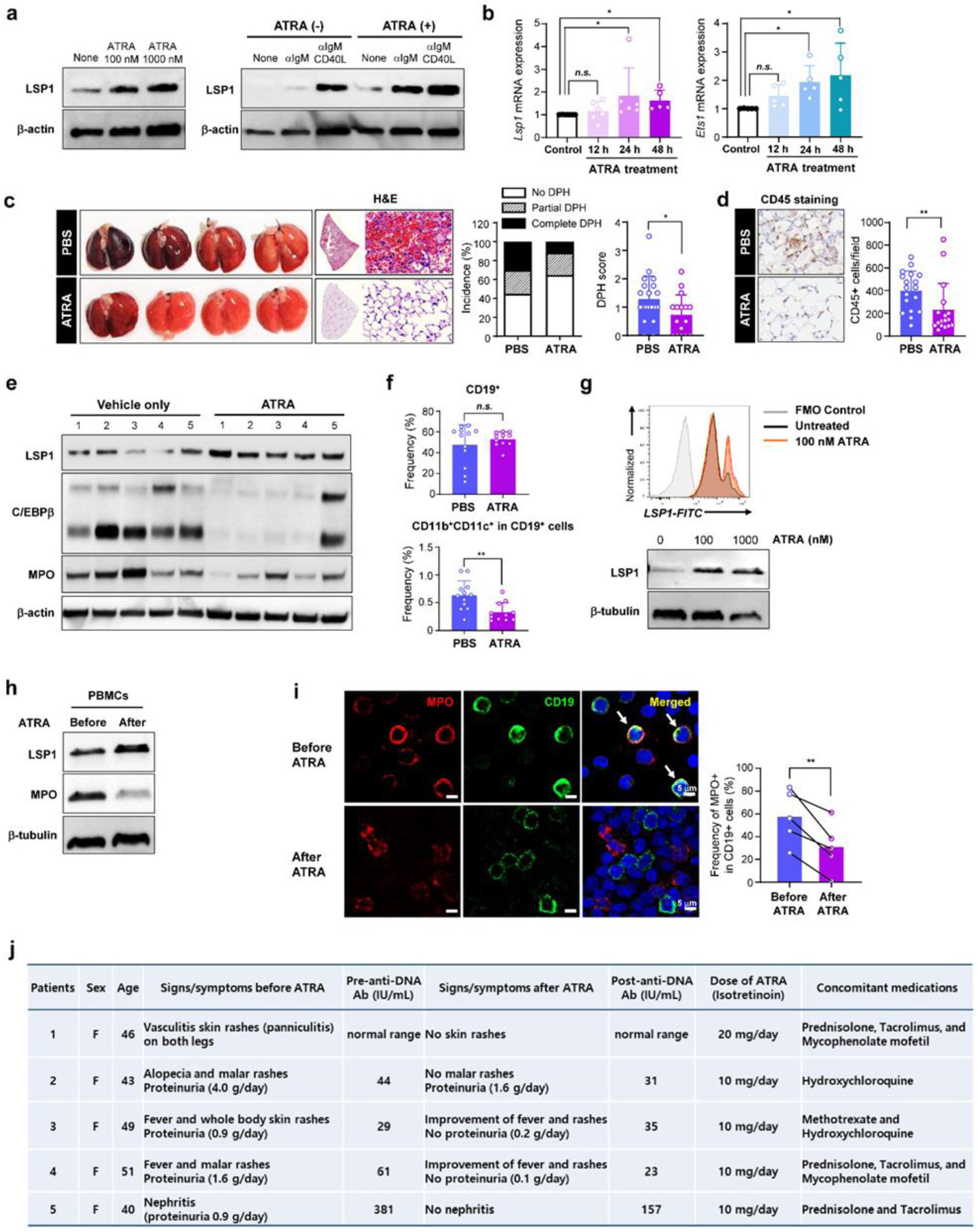
ATRA-mediated suppression of pristane-induced lupus, ABC generation, and myeloperoxidase expression in SLE B cells. **a**, Immunoblotting for LSP1 expression in B cells after ATRA treatment. Splenic B cells were treated with 100 or 1000 nM ATRA for 3 days without stimuli (left panel) or stimulated with αIgM and CD40L in the absence or presence of 100 nM ATRA for 3 days (right panel). **b.** Quantitative RT-PCR analysis of *Lsp1* and *Ets1* mRNA expression in splenic B cells treated with 1000 nM ATRA for indicated time (n=5∼6). **c-f,** Suppressive effects of ATRA on pristane-induced lupus. (**c,d**) One hour before pristane injection, 0.5 mg/kg ATRA (n=17) or PBS (n=19) was administered via the intraperitoneal route to WT mice. Two weeks after pristane injection, lung tissues were harvested and subjected to H&E staining and immunostaining. Lung pathology was evaluated by DPH incidence and score (**c**) and by the extent of infiltration of CD45^+^ leukocytes (**d**), as described in Fig. 5d and 5e. (**e**) Immunoblotting for LSP1 and its target C/EBPβ and MPO in splenic B cells from mice with pristane-induced lupus. The numbers indicate the individual mice. (**f**) Percentages of splenic CD19^+^ B cells and CD11b^+^CD11c^+^ in CD19^+^ cells (ABCs) determined by flow cytometry (n=12 for the PBS group, n=11 for the ATRA group). **g,** ATRA-induced upregulation of LSP1 expression in SLE B cells. After treatment with ATRA for 3 days, LSP1 expression in the B cells was determined by flow cytometry (top) and western blotting (bottom). **h-i,** Downregulation of MPO expression in SLE B cells induced by ATRA treatment. PBMCs were serially obtained from SLE patients before and 3 to 6 months after ATRA treatment. (**h**) LSP1 and MPO expression in PBMCs was examined by immunoblotting. (**i**) Immunofluorescence staining of MPO. Representative images of MPO expression (red) in CD19^+^ B cells (green) are depicted in merged images (white arrows). The numbers of MPO^+^ cells among the CD19^+^ cells were evaluated in 6 fields per sample. Scale bars=5 μm. **j,** Efficacy of ATRA treatment in 5 SLE patients who were unresponsive to conventional therapy. The data in the bar graphs in (**b**-**d**, **f**, and **i**) are presented as the means ± SDs of at least two independent experiments. *P* values were determined by Kruskal-Wallis test with Dunn’s multiple comparison (**b**), Fisher’s exact test (percentage of DPH incidence in **c**), an unpaired two-tailed *t* test (**c, d,** and **f,** except for data on incidence), and a paired *t* test (**i**). *n.s.*, not significant; * *P* < 0.05; ** *P* < 0.01.

In human peripheral B cells, ATRA also upregulated LSP1 expression, as determined by flow cytometry and western blot analysis (**Fig. 7g**), which is consistent with the mouse data. Moreover, LSP1 expression in the peripheral mononuclear cells of drug-naïve SLE patients was upregulated after ATRA treatment for 3 months, accompanied by a dramatic reduction in MPO expression levels (**Fig. 7h**). Concomitantly, after ATRA was administered for 3 to 6 months to 5 SLE patients whose disease was refractory to conventional medications, MPO expression in B cells was markedly downregulated (**Fig. 7i**). Most strikingly, ATRA treatment successfully improved refractory signs and symptoms, including fever, vasculitic rashes, livedo reticularis, and glomerulonephritis, in all 5 SLE patients (**Fig. 7j**).

In summary, our findings suggest that ATRA upregulates LSP1 expression in B cells and attenuates the progression of lupus in both mice and humans, presumably by modulating the LSP1-CEBPβ-MPO axis.

## Discussion

Here, we demonstrated that LSP1, an ABP, is a master regulator of B lymphocyte-mediated innate immune responses. LSP1 is expressed in B cells, and its expression is altered depending on the maturation, development, and activation of B cells. *Lsp1* deficiency markedly upregulates myeloid gene expression in B cells. *Lsp1-*deficient B cells exhibit morphological and functional characteristics of myeloid cells with strong phagocytic activity and high IgG-producing abilities. The PKCβ-CEBPβ pathway is critical for generating chimeric LRBs. *Lsp1* deficiency accelerates the progression of pristane-induced lupus in mice, promoting the myeloid phenotype in B cells and increasing autoantibody production; these changes were reproduced in B-cell-specific *Lsp1* KO mice. Moreover, LSP1 expression is downregulated in SLE patients. LSP1 downregulation can be induced by ETS1 gene defects, lupus-inducing drugs, and TLR7 ligands and is inversely correlated with MPO expression in B cells, disease activity, and the IFN signature in SLE patients. ATRA upregulates LSP1 expression in B cells and attenuates the progression of lupus in both mice and humans possibly through the regulation of the LSP1-CEBPβ-MPO axis.

Emerging evidence has shown that genetic defects related to actin cytoskeleton remodeling, including *WAS*, *NCKAP1L*, *and ARPC1B* defects, are associated with impaired immune function and the development of autoimmune diseases^38^. In actin remodeling, ABPs are essential for maintaining myeloid cell functions, such as phagocytosis, but their role in lymphocytes is poorly understood. According to our interactome analysis, LSP1 was identified as the top-ranked ABP that was expressed in lymphocytes and related to innate immunity. Subsequent transcriptome, morphological, and the *in vitro* functional studies revealed that *Lsp1*-deficient B cells (LRBs) appeared to exhibit a myeloid phenotype. B cells can be converted into myeloid cells via epigenetic modifications when certain transcription factors or signaling transducers defining lineage, including CEBP isoforms, PU.1, and Notch, are overexpressed in B cells^39,40,41^. In contrast to lineage-converted myelocytes from B lymphocytes, LRBs are unique in that they still possess high Ab-producing ability despite their myeloid cell morphology and function. These chimeric attributes seem to facilitate the triggering of autoimmunity by promoting the generation of CD11b^+^CD11c^+^ ABCs and the production of autoantibodies, such as anti-dsDNA Abs.

In recent years, the pathological role of ABCs in aging and autoimmune diseases has gained increased amounts of attention. However, how ABCs are generated has not yet been determined. We demonstrated that CD11b/CD11c expression was upregulated in *Lsp1*^-/-^ B cells. Moreover, the gene expression patterns of LRBs were similar to those of CD11b^+^ and CD11c^+^ B cells. In the pristane- or R848-induced lupus models, the percentages of CD11b^+^ and CD11c^+^ B cells were increased, and these increases were also observed in *Lsp1*^-/-^ mice. Taken together, these findings suggest that LSP1 may function as a major regulator of ABC generation. However, despite their similarity to ABCs, LRBs are different from ABCs in several ways. The most distinctive characteristic of myeloid cells is their ability to produce MPO and ROS in addition to phagocytizing pathogens. In contrast to ABCs, LRBs strongly phagocytose foreign and self-antigens. Like neutrophils, LRBs also secreted high amounts of MPO, exclusively found in myeloid cells and have never been reported in ABCs, and produced high levels of myeloid-specific cytokines (*e.g.,* IL-1β) and ROS. In this regard, compared with ABCs, LRBs can more effectively exacerbate autoimmune inflammation via multiple mechanisms: 1) enhanced autoimmunity to self-antigens, 2) MPO- and ROS-mediated tissue damage, and 3) abundant proinflammatory cytokine production.

Mechanistically, our work identified the LSP1-PKCβ-CEBPβ axis as a novel pathway that drives the chimeric action of LRBs. A previous study reported that ectopic expression of CEBPβ readily induces lineage conversion of B cells to macrophages^39^. CEBP also directly binds to the promoter regions of *Bimp1* and *Irf4* and thereby promotes B-cell activation and differentiation^42^. Here, through master regulator analysis, CEBPβ was strongly suggested to be a key TF regulating the myeloid properties of LRBs, and its expression was high in *Lsp1*^-/-^ B cells. Moreover, CEBPβ siRNA downregulated not only MPO expression but also Bimp1, IRF4, and Xbp1 expression in *Lsp1*^-/-^ B cells. Our results, combined with those of an earlier report^39^, suggest that CEBPβ is a central regulator orchestrating the myeloid phenotype, hyperactivation, and differentiation of B cells under *Lsp1*-deficient conditions (**Fig. 4k**).

Genetic and environmental factors contribute to the complex and multifaceted immune dysregulation observed in patients with SLE. In particular, in both mice and/or humans, downregulated ETS1 expression in T and B lymphocytes has been implicated in the development of lupus^30,43^. In the present study, ETS1 expression correlated positively with LSP1 levels in SLE patients. Mice with pristane-induced lupus exhibited downregulated ETS1 and LSP1 expression in B cells. Moreover, the siRNA experimental results indicated that ETS1 is an upstream regulator of LSP1. Given that ETS1 binds to the LSP1 promoter in B cells^26^, an ETS1 defect may be the major genetic factor underlying the reduced LSP1 expression levels in SLE patients. We presume that in individuals with ETS dysfunction, lupus-inducing drugs (hydralazine and procainamide), epigenetic regulators (5-azacytidine), and TLR7 stimulants (e.g., RNA viruses and endogenous ligands) further exacerbate the downregulation in LSP1 expression to a disease-inducible level.

MPO is an enzyme involved in killing pathogens, such as bacteria, and is mainly produced by neutrophils and activated macrophages. Interestingly, an increase in MPO expression can lead to the excessive production of ROS and the activation of autoimmunity, contributing to inflammation and tissue damage^18^. In the context of lupus, elevated MPO levels are primarily found in neutrophils but are never reported in B cells^44^. Therefore, it was surprising to observe that MPO expression was upregulated in the B cells of SLE patients and pristane-induced lupus model mice. Strikingly, more than 90% of drug-naïve SLE patients, but only < 1% of healthy donors, were positive for MPO in B cells. Based on these findings, we propose that MPO is a single marker for detecting B cells with a myeloid phenotype (probably LRBs) and might be helpful for SLE diagnosis. Furthermore, an MPO inhibitor substantially attenuated the progression of pristane-induced lupus, which suggested that MPO could also be a promising therapeutic target.

Another novel finding of this study was the discovery of ATRA as a new therapeutic agent for upregulating LSP1. Indeed, ATRA has been widely used for the treatment of acne and acute promyelocytic leukemia^45^. Here, ATRA upregulated LSP1 expression in B cells, restored the levels of CEBP/MPO expression and the percentage of CD11b^+^/11c^+^ B cells, and retarded disease progression in mice with lupus. Most importantly, albeit in small-scale clinical trials, ATRA successfully relieved refractory symptoms to conventional drugs in all SLE patients while remarkably downregulating MPO expression in B cells. However, the molecular mechanism through which LSP1 is upregulated by ATRA treatment has not been determined. A study demonstrated that the promoter region of *Ets1* contains a retinoic acid response element (RARE) and that ATRA upregulates *Ets1* expression by binding to RARE in osteoblasts^46^. We observed that ATRA upregulated *Ets1* mRNA in addition to *Lsp1* mRNA in B cells. Although the exact mechanism involved remains elusive, we suspect that ATRA can increase LSP1 expression levels in B cells through ETS1 upregulation.

Overall, the present study provides novel insight into the ’chimeric’ B-cell-mediated acceleration of autoimmune disease. First, our findings suggest a new role for ABPs in lymphocytes, linking LSP1 with the morphological and functional conversion of B cells to myeloid cells. Second, this study identified the novel B-cell subtype LRBs as having both strong phagocytic and high Ab-producing abilities and explained how ABCs or B cells with a myeloid phenotype are generated. Third, we propose an LSP1 signature consisting of 33 genes, including CEBPα, CEBPβ, and MPO, that could be utilized to diagnose SLE and predict its activity. Finally, our translational work highlights the importance of LSP1 in SLE pathogenesis, suggesting that LSP1-elevating drugs, such as ATRA, could be new treatment strategies for SLE. Similarly, the PKCβ-CEBPβ-MPO axis could also be a target for B-cell-dependent autoimmune diseases, as evidenced by the use of an inhibitor of PKCβ (ruboxistaurin) and an MPO inhibitor (PF06281355). We anticipate that our data increases the understanding of the pathogenesis of autoimmune diseases and provides a novel approach that can be applied to diagnose and treat SLE in clinical settings.

## Data and materials availability

### Lead contact

Requests for further information or reagents should be directed to and will be fulfilled by the lead contact, Wan-Uk Kim.

## Acknowledgments

This study was supported by a grant from the National Research Foundation of Korea (NRF) funded by the Ministry of Science and ICT (2015R1A3A2032927 to W.U.K., 2021R1A2C1008130 to N.L., and 2020R1I1A1A01071974 to B.K.H.). We also thank Professor Tae Jin Kim (Sungkyunkwan University School of Medicine, Suwon, Republic of Korea) for critically reviewing this manuscript.

## Author contributions

N.L. and W.U.K. designed the experiments. R.K., J.K., E.C., K.G.L., Y.M.K., Y.L., J.K., Y.J.P., B.K.H., and N.L. performed the experiments. S.Y. and B.K.H. analyzed the NGS data. B.K.H., N.L., and W.U.K. interpreted the results from data analyses. L.S. and S.H.I. kindly provided the *Lsp1^-/-^*and *Ets1^-/-^* mice, respectively. N.L., B.K.H., and W.U.K. wrote and edited the original manuscript. Y.C. and C.S.C provided valuable discussions. All authors commented on the manuscript.

## Materials and Methods

### Prediction of the immunological role of actin-binding proteins

To compile the gene set of actin binding proteins (ABPs), we derived a list of genes categorized under the “actin binding” term (GO:0003779) in the GOMF category in the Gene Ontology database (https://www.geneontology.org/). The protein expression dataset was extracted from the ‘Gene’, ‘Cell type’, and ‘Level’ columns after downloading ’Normal tissue data’ stored in the Human Protein Atlas (https://www.proteinatlas.org/), which represents the most comprehensive database for the spatial distribution of human proteins in tissues and cells. To define ABPs expressed in lymphocytes, macrophages, and hematopoietic cells, we filtered the protein dataset according to cell type and selected the proteins that were included in the list of ABPs we identified. Proteins with a ’level of low’, which indicates a low degree of expression, were excluded. Next, we examined the protein‒protein interaction network, which was constructed using the STRING database (https://string-db.org/), of the curated ABPs and identified the first neighbors that interacted with each ABP. Finally, we conducted an enrichment analysis of the complete gene sets of ABPs using the ’enrichGO’ function from the ’clusterProfiler’ package in R (version 4.2.1).^1^ Finally, we summarized and visualized the resulting GOBP terms with REVIGO (http://revigo.irb.hr/).^2^

### Mice

All of the mice used in this study were of the C57BL/6 genetic background and were maintained under specific pathogen-free conditions. WT mice were used as age- and sex-matched controls. Mice genetically deficient in the *Lsp1* gene (*Lsp1^-/-^*) were kindly provided by Dr. Laurent Sabbagh (Domain Therapeutics North America Inc., Montréal, Québec, Canada).^3^ To generate B-cell-specific *Lsp1^-/-^* mice, hereafter termed *Lsp1^fl/fl^ Cd19^Cre/-^* mice, we used the *loxP/Cre* system, and *Lsp1*^flox/flox^ (*Lsp1^fl/fl^*) mice were used as control mice. The genotyping of these mice was performed by PCR analysis of the *flox* and *Cd19 Cre* genes using the following primer sequences: forward for the *flox* gene, 5’-CCCTAGGTCTCTTACATCACAGC-3’; reverse for the *flox* gene, 5’-ACTGGTTAGACAGAGGATGTTGG -3’; forward for the *Cd19 Cre* gene, 5’-GCGGTCTGGCA GTAAAAACTATC-3’; reverse for the *Cd19 Cre* gene, 5’-GTGAAACAGCATTGCTGTCACTT-3’. WT (or *Ets1^fl/fl^*) and *Ets1^-/-^* (or *Ets1^fl/fl^Cd19^Cre/-^*) mice were generously provided by Dr. Sin-Hyeog Im (Pohang University of Science and Technology, Pohang, Korea). The animal experimental procedures were approved by the Institutional Animal Care and Use Committee (IACUC) of the Catholic University of Korea and performed in accordance with the guidelines and policies provided by the IACUC.

### B-cell isolation and activation

Human and mouse B cells were isolated from human peripheral blood and mouse splenocytes using a human B-cell isolation kit II (Miltenyi Biotech, #130-091-151) and a mouse pan-B-cell isolation kit II (Miltenyi Biotech, #130-104-443), respectively, according to the manufacturer’s protocol. The cells were cultured in RPMI-1640 media (Welgene, #LM-011-03) supplemented with 10% fetal bovine serum (FBS; Gibco, #16000044), 1% antibiotics (Gibco, #15140-122), and 1 mM sodium pyruvate (Gibco, #11360-070). B cells were activated with an anti-human or anti-mouse IgM antibody (10 μg/mL each; Jackson ImmunoResearch, #309-006-043 for human, #115-006-020 for mouse), recombinant human or mouse CD40 ligand (0.1 μg/mL; Enzo, #ALX-522-110 for human, #ALX-522-120 for mouse), lipopolysaccharides (LPS) from *E. coli* O111:B4 (2 μg/mL; Sigma, #L2630), CpG-ODN2006 (1 μg/mL; InvivoGen, #tlrl-2006), and R848 (1 μg/mL; InvivoGen, #tlrl-r848). In some experiments, recombinant HVEM (5 μg/mL; Enzo, #ALX-522-017-C050), PD-1 (5 μg/mL; Enzo, #ENZ-PRT190-0050), BAPTA (5 μM; Invitrogen, #B1205), the PKCβ inhibitor ruboxistaurin (4 μM; MedChemExpress, #HY-10195), procainamide (20 μM; Sigma‒Aldrich, #P9391), hydralazine (20 μM; Sigma‒Aldrich, #H1753), 5-azacytidine (1 μM; Sigma‒Aldrich, #A2385), or ATRA (100 and 1000 nM; Sigma‒Aldrich, #R2625) was added to B cells as indicated. For gene knockdown, human *LSP1* siRNA (Santa Cruz Biotechnology, #sc-42899), *ETS1* siRNA (Santa Cruz Biotechnology #sc-29309), or mouse *PKCβ* siRNA (Santa Cruz Biotechnology, #sc-36255) was electroporated into B cells using an Amaxa Nucleofector (Lonza) and incubated as indicated. In the case of CEBPβ knockdown, *Lsp1^-/-^* B cells were pretreated with 5 μM polybrene B (Santa Cruz Biotechnology, #sc-134220) and transduced with control or CEBPβ shRNA-containing lentiviral particles (Santa Cruz Biotechnology, #sc-108080, #sc-29862-v).

### Bulk RNA sequencing analysis and data processing

B cells isolated from wild-type (WT) and *Lsp1^-/-^* mice were treated with media alone, αIgM, or LPS for 4 hours, after which total RNA was extracted using an RNeasy Mini Kit (QIAGEN, #74106) according to the manufacturer’s instructions. Bulk RNA sequencing analysis was performed using 1 μg of RNA depleted of ribosomal RNA. At least 59.15 M reads (101 bp paired-end) were sequenced. The quality control pipeline utilized the Illumina standards to assess base call quality and truncate low-quality reads. Sequence alignment and quantification were performed using the STAR-RSEM pipeline.^4,5^ Reads overlapping exons in the annotation of the latest version of the NCBI RefSeq database were identified. Reads overlapping exons in the annotation of the Genome Reference Consortium Human Build 38 (GRCh38) were identified. Genes were filtered out and excluded from downstream analysis if they failed to achieve a raw read count of at least 2 across all the libraries. Batch effects were corrected by ComBat.^6^ The trimmed mean of M-values normalization method (TMM) (version 1.6.1)^7^ was used for calculating normalized count data.

### Differential expression analysis

The integrated hypothesis testing method^7^ was applied to determine the DEGs between WT and *Lsp1*-deficient B cells. Briefly, t tests and median difference tests were performed for each gene, and the *P* values from the two tests were integrated into one combined *P* value using Stouffer’s method.^8^ The false discovery rate (FDR) was computed from the combined *P* value using Story’s correction method.^9^ DEGs with an FDR<0.05 and a log2-fold change≥1 were selected, which was the value at the 95th percentile of the log2-fold change distribution from the RNA-sequencing data. A list of DEGs associated with *Lsp1* deficiency was generated. Functional enrichment analysis was conducted using DAVID software^10^ to identify the enriched cellular processes and molecular pathways were associated with the DEGs.

### Quantitative real-time PCR (qRT‒PCR)

For qRT‒PCR, cDNA was synthesized using RevertAid Reverse Transcriptase (Thermo Fisher Scientific, #0441), and PCR was performed in a CFX96 real-time PCR system using iQ SYBR Green Supermix (Bio-Rad, #1708880). The primers used for qRT‒PCR are listed in **Extended Data Table 2**. All samples were normalized to *Gapdh* expression, and relative mRNA expression levels or fold changes were calculated using the *2^−ΔΔCt^* method.

### Immunoblotting

Human and murine B cells were lysed in RIPA buffer (Thermo Fisher, #89900) supplemented with protease inhibitors (Roche, #11697498001). Protein concentrations were determined using a BCA assay kit (Pierce, #23225). Equal amounts of proteins were separated via SDS‒PAGE and subsequently transferred onto PVDF membranes using a Trans-blot turbo transfer system (all from Bio-Rad, #1704156). The primary Abs used were as follows: LSP1 (BD Biosciences for human; #610734; and Cell Signaling Technology for murine cells; #3812, respectively); MPO (R&D Systems; #AF3667); Blimp1 (Santa Cruz Biotechnology; #sc-47732); Xbp1 (Santa Cruz Biotechnology; #sc-8015); IRF4 (Santa Cruz Biotechnology; #sc-48338); C/EBPβ (Abcam; #ab32358); PKCβ (Invitrogen; #PA5-64496); PKCβ (GeneTex; #113252); IL-1β (R&D Systems; #AF401-NA); ETS1 (Cell Signaling Technology; #14069); GAPDH (Santa Cruz Biotechnology; #sc-25778); β-actin (Santa Cruz Biotechnology; #sc-47778); and β-tubulin (Abcam; #15568). Horseradish peroxidase-conjugated anti-rabbit IgG (Thermo Fisher Scientific, #31460), anti-goat IgG (Santa Cruz Biotechnology, #sc-2020), and anti-mouse IgG (Thermo Fisher Scientific, #31430) were used as secondary Abs. The signals on the membranes were visualized using a chemiluminescent detection system (Pierce, #32106, #34095).

### Flow cytometry

Single-cell suspensions were prepared from the spleens, lymph nodes, and BALF of WT and *Lsp1^-/-^* mice, as well as from human peripheral blood mononuclear cells (PBMCs). To block nonspecific Ab binding, cells were pretreated with an FcR blocker (Miltenyi Biotechnology, #130-092-575 for mouse; BioLegend, #422302 for human) for 10 minutes. Surface staining was carried out for 30 minutes at 4°C with the following fluorochrome-labeled anti-mouse Abs: APC/Cy7-CD45 (BD Bioscience, #557659), PE/Cy5-B220 (eBioscience, #15-0452-82), FITC-GL7 (BioLegend, #144604), PE-CD95 (BioLegend, #152608), APC-CD138 (Invitrogen, #142506), eFluor660-IgM (eBioscience, #50-5790-82), PE-IgD (eBioscience, #12-5993-82), APC/Cy7-CD21 (BioLegend, #123418), PE/Cy7-CD23 (eBioscience, #25-0232-82), PE/Cy7-CD19 (Invitrogen, #25-0193-82), Brilliant Violet 510-CD11b (BioLegend, #101263), PE/Cy5-CD11c (BioLegend, #117316), and PE-CSF1R (Invitrogen, #12-1152-82). The following fluorochrome-labeled anti-human Abs were used for surface staining: PerCP/Cy5.5-CD19 (Invitrogen, #45-0198-42), APC-CD27 (eBioscience, #17-0279-42), Brilliant Violet 421-IgD (BioLegend, #348226), and PE-MPO (Invitrogen, #12-1299-41). The intracellular expression of LSP1 in B cells was also detected by flow cytometry as described previously.^11^ In brief, the cells were fixed, permeabilized, and incubated with rabbit anti-LSP1 Ab (Cell Signaling Technology, #3812 for mouse) or recombinant rabbit IgG (Abcam, #ab172730) for 1 hour and then stained with a FITC-conjugated anti-rabbit Ab (Invitrogen, #31635) for 30 minutes to determine LSP1 expression. For human B cells, the cells were stained with a FITC-conjugated mouse anti-LSP1 Ab (BD Biosciences, #610734 for human) for 1 hour. FITC-conjugated mouse IgG_1_ (Santa Cruz Biotechnology, #sc-3877) was used as an isotype control. The cells were resuspended in 1% bovine serum albumin (BSA) containing PBS and analyzed with a FACSCanto II (BD Biosciences) or an LSR Fortessa (BD Biosciences) with DIVA software. All of the data were analyzed using FlowJo software (FlowJo).

### Immunohistochemistry and immunofluorescence

For immunohistochemistry, mouse lung tissues were fixed in 4% paraformaldehyde and embedded in paraffin. The paraffin sections were deparaffinized in xylene, followed by antigen retrieval in sodium citrate buffer. After the removal of endogenous peroxidases with 3% hydrogen peroxide for 30 minutes, the sections were blocked with 10% normal donkey serum (Jackson ImmunoResearch, #017-000-121) and stained with Abs against CD45 (Abcam, #ab64100) or IgG (Abcam, #ab197767) overnight at 4°C. As a secondary Ab, goat anti-rat IgG-horseradish peroxidase (Vector Laboratory, #MP-7404) or horse anti-goat IgG-horseradish peroxidase (Vector Laboratory, #MP-7405) was then reacted with the sections for 2 hours at room temperature. The tissues were incubated with 3,3’-diaminobenzidine (DAB) (Vector Laboratories, #SK-4100) for visualization, and the nuclei were counterstained with Mayer’s hematoxylin solution (Sigma‒Aldrich, #MHS32). Images were taken with a Panoramic MIDI slide scanner (3DHISTECH).

For immunofluorescence staining, human PBMCs or murine B cells were fixed in 4% paraformaldehyde (Wako Pure Chemicals, #16320145), mounted on slides, and blocked with 10% normal donkey serum. Alexa Fluor 488-conjugated phalloidin (Invitrogen, #A12379), B220 (Abcam, #10558), MPO (R&D Systems, #AF3667), and CD19 (Abcam, #134114) Abs were added and incubated overnight at 4°C. The sections were subsequently incubated with chicken anti-rat IgG Alexa Fluor 647 (Invitrogen, #A-21472), donkey anti-rabbit IgG Alexa 488 (Invitrogen, #A-21206), or donkey anti-goat IgG Alexa Fluor 594 (Invitrogen, #A11058) secondary Abs for 1 hour at room temperature. The cells were stained with 4′,6-diamidine-2′-phenylindole (DAPI; Roche, #10236276001). Images were taken with a Zeiss LSM 900 microscope (Carl Zeiss) and contrast-enhanced using Zeiss ZEN microscope software.

### Enzyme-linked immunosorbent assay (ELISA)

The levels of MPO, TNF-α, and IgG in the culture supernatants from stimulated B cells were determined by ELISA kits for human MPO (R&D Systems, #DY3174), mouse MPO (R&D Systems, #DY3667), mouse TNF-α (R&D Systems, #DY410), and human IgG (eBioscience, #88-50550) according to the manufacturers’ instructions. Anti-dsDNA IgG levels in the sera of mice with pristane-induced lupus were measured by an ELISA kit (Alpha Diagnostics, #5120) according to the manufacturer’s instructions. To detect total IgG, IgG2c, IgG3, and IgM levels, ELISA was also performed. Briefly, goat anti-mouse IgG (all Abs from Southern Biotech, #1030-01), goat anti-mouse IgG2c (#1070-01), goat anti-mouse IgG3 (#1100-01), and goat anti-mouse IgM (#1020-01) in PBS were added to 96-well EIA plates (Costar, #9018), which were incubated overnight at 4°C and then blocked with 1% BSA in PBS for 2 hours at room temperature. The samples were incubated for 2 hours, after which they were incubated with horseradish peroxidase-conjugated donkey anti-mouse IgG (H+L; #6410-15) as a detection Ab for an additional 2 hours at room temperature. Finally, the samples were developed with 3,3′,5,5′-tetramethylbenzidine (TMB; R&D Systems, #DY999) solution, and the reaction was stopped with 2% H_2_SO_4_. The absorbance at 450 nm (OD_450nm_) was measured by an ELISA reader (Perkin Elmer).

### Efferocytosis

Isolated thymocytes were treated with 1.0 μM dexamethasone (Sigma‒Aldrich, #D2915) in complete RPMI for 18 hours, after which apoptotic cells were assessed via FITC-Annexin V staining (BD Bioscience, #556547) and flow cytometry analysis. Apoptotic thymocytes were labeled with 1 μg/mL pHrodo Red succinimidyl ester (pHrodo Red-SE, Thermo Fisher, #P36600) according to the manufacturer’s instructions and coincubated with stimulated splenic B cells for 30 minutes. B cells engulfing apoptotic thymocytes were analyzed using confocal microscopy (LSM900, Zeiss).

### Live-cell imaging analysis for phagocytosis

WT and *Lsp1^-/-^* B cells stimulated with αIgM were treated with Alexa Fluor 594-conjugated *E. coli* bioparticles (Invitrogen, #E23370) at a ratio of 10 *E. coli* particles per B cell, after which live cell imaging analysis was carried out for 30 minutes at a rate of 1 frame per 30 or 40 seconds using BioTek LION Heart XL (Agilent) or TomoStudio (Tomocube, Inc.). Time-lapse images were acquired in a bright field at 4x magnification using an INCell 2200 image analyzer or TomoAnalysis software.

### Master regulator analysis of DEGs in Lsp1^-/-^ B cells

Master regulator analysis (MRA) was carried out to identify key transcriptional regulators that drive the differential expression patterns observed in the control versus *Lsp1*-deficienct B cells. Data on TFs and their target interactions were collected from publicly available databases, including the Transcriptional Regulatory Element Database (TRED)^12^, Immune Epitope Database (IEDB)^13^, Molecular Signature Database (MSigDB)^14^, Amadeus^15^, bZIPDB^16^, and OregAnno^17^. The targets of individual transcription factors were counted among the DEG. To compute the statistical significance of the number of TF target genes, the following procedures were conducted: 1) the same number of genes as the number of DEGs was randomly sampled from the whole genome, 2) the target genes of each TF were counted in the randomly sampled genes, 3) this procedure was repeated 100,000 times to generate an empirical null distribution, and 4) the significance of an observed target count in the DEGs was computed using a one-tailed test with an empirical null distribution. The TFs whose targets were significantly enriched (*P* < 0.05) by the DEGs were selected as key TFs. The protein‒ DNA interaction network between the transcriptional regulators and targets was visualized using Cytoscape.

### Analysis of public gene expression data

To characterize the transcriptional changes in *Lsp1^-/-^* B cells in association with ABCs, a Gene Set Enrichment Analysis (GSEA) was carried out. To this end, we created a custom gene set consisting of 201 DEGs in cluster 1 that were commonly upregulated in *Lsp1^-/-^* B cells stimulated with media, aIgM, or LPS (*see* **Extended Data Fig. 4a**). The molecular profiling data for GSEA were sourced from the three independent transcriptome datasets as follows: CD11b^+^ B cells (GSE29717), CD11c^+^ B cells (GSE110999), and age-associated B cells (GSE28887).

### KLH immunization and pristane-induced lupus establishment in mice

For KLH immunization, mice were subcutaneously injected with 100 μg of KLH (Sigma‒Aldrich, #H8283) emulsified in complete Freund’s adjuvant (CFA; Sigma‒Aldrich, #F5881) and sacrificed on day 7. For the pristane-induced lupus model, mice were intraperitoneally injected with 500 μl of pristane (Sigma‒Aldrich, #P9622). After 11 to 14 days, lung tissues isolated from the mice were fixed with 4% paraformaldehyde and embedded in paraffin. The paraffin sections were deparaffinized and stained with EASYSTAIN Harris hematoxylin (YD Diagnostics, #S2-5) and eosin Y (Showa, #0501-5330). BALF samples were obtained from the lung tissues using PBS and subjected to flow cytometry analysis. The pathological severity in lungs and BALFs of WT and *Lsp1^-/-^* mice were initially evaluated via gross inspection. The incidence and severity of diffuse pulmonary hemorrhage (DPH) were also determined by H&E staining based on the following scoring system: 0, no hemorrhage; 1, 0–25%; 3, 25–50%; 3, 50–75%; and 4, 75–100%.^18^

### Isolation of PBMCs from SLE patients

This study was approved by the Institutional Review Board of The Catholic University of Korea (IRB approval no. KC15TISI0753). Human peripheral blood was obtained from sex- and age-matched healthy donors and SLE patients who provided informed consent. Peripheral blood mononuclear cells (PBMCs) were isolated by Ficoll-Paque (GE Healthcare, #GE17-1440-02) density gradient centrifugation. SLE disease severity was assessed according to the criteria based on the SLEDAI^19^ as follows: 1∼4, mild; 5∼8, moderate; and 9 or more, severe.

### Assessment of the LSP1 signature in PBMCs of SLE patients

We arbitrarily defined 33 genes, namely, CEBPα, CEBPβ, and 31 target genes of CEBP, in *Lsp1*-deficient B cells as the **‘**LSP1 signature’ (**Fig. 4b**). To evaluate the LSP1 signature in PBMCs from patients with SLE, we analyzed the gene expression profiles obtained from the following public databases: GSE72509, which contains bulk RNA-sequencing data, and GSE174188, which includes multiplexed single-cell RNA-sequencing data on approximately 1.2 million PBMCs. To investigate the LSP1 signature across datasets, we computed Z scores for the 33-gene set for each sample. The Z score calculations for single-cell RNA-seq data were performed using the ’tl.score_genes’ function in Scanpy.^20^ Additionally, to infer the immune cell composition in the PBMCs of SLE patients, bulk RNA-seq data were subjected to deconvolution analysis using the ’MCP-counter’ algorithm, which allows us to estimate the abundance of specific cell types within the heterogeneous PBMC population.^21^

### Selection of candidate chemicals that upregulate LSP1 expression

The Comparative Toxicogenomics Database (CTD) is a publicly available database designed to enhance the understanding of the effects of environmental exposures on human health.^22^ The CTD provides curated information on associations and inferences between chemicals and diseases. In this study, we used the chemical–gene interaction query of CTD to identify chemicals that have an effect on LSP1 expression. From these data, we compiled a dataset of chemicals known to upregulate the expression or activity of LSP1.

### Statistical analysis

Statistical significance was assessed by the unpaired, paired two-tailed Student’s *t* test, Mann‒Whitney *U*-test, one-way, two-way analysis of variance (ANOVA), Wilcoxon two-tailed rank sum test, or Fisher’s exact test, as appropriate. Pearson’s correlation was used to assess the association between two variables. MATLAB (v.9.0; MathWorks, Natick, MA, USA), R (v.3.5), and Python (v.3.7) were used for bioinformatics analysis. A *P* value less than 0.05 was considered statistically significant.

## References

1. Cancro, M.P. Age-Associated B Cells. Annu Rev Immunol 38, 315–340 (2020).

2. Riley, R.L., Khomtchouk, K. & Blomberg, B.B. Age-associated B cells (ABC) inhibit B lymphopoiesis and alter antibody repertoires in old age. Cell Immunol 321, 61–67 (2017).

3. Wang, S. et al. IL-21 drives expansion and plasma cell differentiation of autoreactive CD11c(hi)T-bet(+) B cells in SLE. Nat Commun 9, 1758 (2018).

4. Dorshkind, K., Höfer, T., Montecino-Rodriguez, E., Pioli, P.D. & Rodewald, H.R. Do haematopoietic stem cells age? Nat Rev Immunol 20, 196–202 (2020).

5. Lassar, A.B., Paterson, B.M. & Weintraub, H. Transfection of a DNA locus that mediates the conversion of 10T1/2 fibroblasts to myoblasts. Cell 47, 649–656 (1986).

6. Record, J., Saeed, M.B., Venit, T., Percipalle, P. & Westerberg, L.S. Journey to the Center of the Cell: Cytoplasmic and Nuclear Actin in Immune Cell Functions. Front Cell Dev Biol 9, 682294 (2021).

7. Haynes, J., Srivastava, J., Madson, N., Wittmann, T. & Barber, D.L. Dynamic actin remodeling during epithelial-mesenchymal transition depends on increased moesin expression. Mol Biol Cell 22, 4750–4764 (2011).

8. Le, N.P., Channabasappa, S., Hossain, M., Liu, L. & Singh, B. Leukocyte-specific protein 1 regulates neutrophil recruitment in acute lung inflammation. Am J Physiol Lung Cell Mol Physiol 309, L995–1008 (2015).

9. Kwon, R. et al. Regulation of tumor growth by leukocyte-specific protein 1 in T cells. J Immunother Cancer 8 (2020).

10. Hwang, S.H. et al. Leukocyte-specific protein 1 regulates T-cell migration in rheumatoid arthritis. Proc Natl Acad Sci U S A 112, E6535–6543 (2015).

11. Wu, J.L. et al. Temporal regulation of Lsp1 O-GlcNAcylation and phosphorylation during apoptosis of activated B cells. Nat Commun 7, 12526 (2016).

12. Swaminathan, A., Lucas, R.M., Dear, K. & McMichael, A.J. Keyhole limpet haemocyanin - a model antigen for human immunotoxicological studies. Br J Clin Pharmacol 78, 1135–1142 (2014).

13. Klein, D.P., Jongstra-Bilen, J., Ogryzlo, K., Chong, R. & Jongstra, J. Lymphocyte-specific Ca2+-binding protein LSP1 is associated with the cytoplasmic face of the plasma membrane. Mol Cell Biol 9, 3043–3048 (1989).

14. Pulford, K., Jones, M., Banham, A.H., Haralambieva, E. & Mason, D.Y. Lymphocyte-specific protein 1: a specific marker of human leucocytes. Immunology 96, 262–271 (1999).

15. Klaus, G.G., Choi, M.S. & Holman, M. Properties of mouse CD40. ligation of CD40 activates B cells via a Ca(++)-dependent, FK506-sensitive pathway. Eur J Immunol 24, 3229–3232 (1994).

16. Chung, M.K.Y. et al. Functions of double-negative B cells in autoimmune diseases, infections, and cancers. EMBO Mol Med 15, e17341 (2023).

17. Di Tullio, A. et al. CCAAT/enhancer binding protein alpha (C/EBP(alpha))-induced transdifferentiation of pre-B cells into macrophages involves no overt retrodifferentiation. Proc Natl Acad Sci U S A 108, 17016–17021 (2011).

18. Davies, M.J. & Hawkins, C.L. The Role of Myeloperoxidase in Biomolecule Modification, Chronic Inflammation, and Disease. Antioxid Redox Signal 32, 957–981 (2020).

19. Tarazona-Santos, E. et al. Evolutionary dynamics of the human NADPH oxidase genes CYBB, CYBA, NCF2, and NCF4: functional implications. Mol Biol Evol 30, 2157–2167 (2013).

20. Nutt, S.L., Fairfax, K.A. & Kallies, A. BLIMP1 guides the fate of effector B and T cells. Nat Rev Immunol 7, 923–927 (2007).

21. Griffin, D.O. & Rothstein, T.L. A small CD11b(+) human B1 cell subpopulation stimulates T cells and is expanded in lupus. J Exp Med 208, 2591–2598 (2011).

22. Rubtsov, A.V. et al. Toll-like receptor 7 (TLR7)–driven accumulation of a novel CD11c+ B-cell population is important for the development of autoimmunity. Blood 118, 1305–1315 (2011).

23. Torres-Avilés, N.A. et al. Exposure to p,p’-DDE Induces Morphological Changes and Activation of the PKCα-p38-C/EBPβ Pathway in Human Promyelocytic HL-60 Cells. Biomed Res Int 2016, 1375606 (2016).

24. Zhuang, H. et al. Pathogenesis of Diffuse Alveolar Hemorrhage in Murine Lupus. Arthritis Rheumatol 69, 1280–1293 (2017).

25. Reeves, W.H., Lee, P.Y., Weinstein, J.S., Satoh, M. & Lu, L. Induction of autoimmunity by pristane and other naturally occurring hydrocarbons. Trends Immunol 30, 455–464 (2009).

26. Omori, S.A., Smale, S., O’Shea-Greenfield, A. & Wall, R. Differential interaction of nuclear factors with the leukocyte-specific pp52 promoter in B and T cells. J Immunol 159, 1800–1808 (1997).

27. Nickerson, K.M. et al. Age-associated B cells are heterogeneous and dynamic drivers of autoimmunity in mice. J Exp Med 220, e20221346 (2023).

28. Gladman, D.D., Ibañez, D. & Urowitz, M.B. Systemic lupus erythematosus disease activity index 2000. J Rheumatol 29, 288–291 (2002).

29. Tsokos, G.C. Systemic lupus erythematosus. N Engl J Med 365, 2110–2121 (2011).

30. Garrett-Sinha, L.A., Kearly, A. & Satterthwaite, A.B. The Role of the Transcription Factor Ets1 in Lupus and Other Autoimmune Diseases. Crit Rev Immunol 36, 485–510 (2016).

31. Brown, G.J. et al. TLR7 gain-of-function genetic variation causes human lupus. Nature 605, 349–356 (2022).

32. He, Y. & Sawalha, A.H. Drug-induced lupus erythematosus: an update on drugs and mechanisms. Curr Opin Rheumatol 30, 490–497 (2018).

33. Borchers, A.T., Keen, C.L. & Gershwin, M.E. Drug-induced lupus. Ann N Y Acad Sci 1108, 166–182 (2007).

34. Stresemann, C. & Lyko, F. Modes of action of the DNA methyltransferase inhibitors azacytidine and decitabine. Int J Cancer 123, 8–13 (2008).

35. Hung, T. et al. The Ro60 autoantigen binds endogenous retroelements and regulates inflammatory gene expression. Science 350, 455–459 (2015).

36. Perez, R.K. et al. Single-cell RNA-seq reveals cell type-specific molecular and genetic associations to lupus. Science 376, eabf1970 (2022).

37. Psarras, A., Wittmann, M. & Vital, E.M. Emerging concepts of type I interferons in SLE pathogenesis and therapy. Nat Rev Rheumatol 18, 575–590 (2022).

38. Dupré, L. & Prunier, G. Deciphering actin remodelling in immune cells through the prism of actin-related inborn errors of immunity. Eur J Cell Biol 102, 151283 (2023).

39. Cirovic, B. et al. C/EBP-Induced Transdifferentiation Reveals Granulocyte-Macrophage Precursor-like Plasticity of B Cells. Stem Cell Reports 8, 346–359 (2017).

40. Heinz, S. et al. Simple combinations of lineage-determining transcription factors prime cis-regulatory elements required for macrophage and B cell identities. Mol Cell 38, 576–589 (2010).

41. Xiu, Y. et al. Coactivation of NF-κB and Notch signaling is sufficient to induce B-cell transformation and enables B-myeloid conversion. Blood 135, 108–120 (2020).

42. Pal, R. et al. C/EBPbeta regulates transcription factors critical for proliferation and survival of multiple myeloma cells. Blood 114, 3890–3898 (2009).

43. Sunshine, A. et al. Ets1 Controls the Development of B Cell Autoimmune Responses in a Cell-Intrinsic Manner. Immunohorizons 3, 331–340 (2019).

44. Telles, R.W., Ferreira, G.A., da Silva, N.P. & Sato, E.I. Increased plasma myeloperoxidase levels in systemic lupus erythematosus. Rheumatol Int 30, 779–784 (2010).

45. Wang, Z.Y. & Chen, Z. Acute promyelocytic leukemia: from highly fatal to highly curable. Blood 111, 2505–2515 (2008).

46. Raouf, A., Li, V., Kola, I., Watson, D.K. & Seth, A. The Ets1 proto-oncogene is upregulated by retinoic acid: characterization of a functional retinoic acid response element in the Ets1 promoter. Oncogene 19, 1969–1974 (2000).

