## Supplementary information for "B lymphocytes acquire myeloid and autoimmune phenotypes via the downregulation of lymphocyte-specific protein-1"

[1]Center for Integrative Rheumatoid Transcriptomics and Dynamics, The Catholic University of Korea, Seoul, Republic of Korea
[2]Division of Cancer Biology and Therapeutics, Department of Surgery, Samuel Oschin Comprehensive Cancer Institute, Cedars-Sinai Medical Center, Los Angeles, CA 90048; Division of Cancer Biology and Therapeutics, Department of Biomedical Sciences, Samuel Oschin Comprehensive Cancer Institute, Cedars-Sinai Medical Center, Los Angeles, CA 90048; Division of Cancer Biology and Therapeutics, Department of Pathology and Laboratory Medicine, Samuel Oschin Comprehensive Cancer Institute, Cedars-Sinai Medical Center, Los Angeles, CA 90048, USA
[3]Lab of Immune Regulation, Research Institute of Pharmaceutical Sciences, College of Pharmacy, Seoul National University, Seoul, Republic of Korea
[4]Department of Pharmacy, Jeju National University, Jeju, Republic of Korea
[5]Department of Life Sciences, Pohang University of Science and Technology (POSTECH), Pohang, Republic of Korea
[6]Domain Therapeutics North America Inc., Montréal, Québec, Canada
[7]Division of Rheumatology, Department of Internal Medicine, The Catholic University of Korea, Seoul, Republic of Korea
[8]Department of Biomedicine & Health Sciences, The Catholic University of Korea, Seoul, Republic of Korea

[*]Both Naeun Lee and Bong-Ki Hong are equally contributed as co-first authors.

Correspondence and reprint request to:

Dr. Wan-Uk Kim, Division of Rheumatology, Department of Internal Medicine, Catholic University of Korea, School of Medicine, Seoul, Korea

**Keyword:** Lymphocyte-specific protein-1 (LSP1), actin-binding proteins (ABPs), B lymphocytes, LSP1-CEBPβ-MPO axis, myeloidization

**This file includes:**

**Extended Data Table 1.** The top 10 actin-binding proteins (ABPs) expressed in lymphocytes, macrophages, and hematopoietic cells according to their correlation with the GOBP term "immune response".

**Extended Data Table 2.** Primer sequences for qRT–PCR

**Extended Data Figure 1.** Identification of actin-binding protein (ABP)s related to immune responses in lymphocytes.

**Extended Data Figure 2.** Effect of LSP1 on the polarization of T helper cells and the generation of plasma cells in the KLH immunization model.

**Extended Data Figure 3.** Upregulation of LSP1 expression in murine B cells by costimulatory signals but not by coinhibitory signals.

**Extended Data Figure 4.** Gene expression profile of splenic B cells from *Lsp1*$^{-/-}$ mice.

**Extended Data Figure 5.** LSP1 regulation of the genes involved in B-cell differentiation and myeloid cell activation.

**Extended Data Figure 6.** Effect of *Lsp1* knockout on the frequency of MPO$^{+}$ B cells and the production of IgM.

**Extended Data Figure 7.** No effect of *Lsp1* knockout on B-cell survival and proliferation.

**Extended Data Figure 8.** Ultrastructural analysis of WT and *Lsp1*$^{-/-}$ B cells.

**Extended Data Figure 9.** No change in C/EBPβ expression in *Lsp1*$^{-/-}$ B cells by ERK inhibition.

**Extended Data Figure 10.** Comparison of the percentages of B cells and myeloid cells in the BALFs of WT and *Lsp1*$^{-/-}$ mice injected with pristane.

**Extended Data Figure 11.** LSP1 expression in diverse cell types from B-cell-specific *Lsp1*-knockout mice.

**Extended Data Figure 12.** Effect of *LSP1* deficiency on ETS1 expression in human B cells.

**Extended Data Figure 13.** Changes in the expression of LSP1 and myeloid genes in B cells following repetitive exposure to the TLR7 agonist R848.

**Extended Data Figure 14.** TEM images of healthy and SLE B cells.

**Extended Data Figure 15**. Amelioration of pristane-induced lupus by an MPO inhibitor.

**Extended Data Figure 16.** The bar graph shows the top 10 compounds that can upregulate LSP1 expression

**Extended Data Video 1 and 2.** Phagocytic activity of WT (Video 1, left panel) and *Lsp1*$^{-/-}$ B cells (Video 2, right panel).

**Extended Data Video 3.** TomoCube images showing the phagocytic activity of *Lsp1*$^{-/-}$ B cells.

**Extended Data Video 4.** Inhibition of the phagocytic activity of *Lsp1*$^{-/-}$ B cells by an actin polymerization inhibitor.

**Extended Data References**

**Extended Data Methods**

**Extended Data Table 1.** The top 10 actin-binding proteins (ABPs) expressed in lymphocytes, macrophages, and hematopoietic cells according to their correlation with the GOBP term "immune response".

| Rank | Lymphocyte | | | Macrophage | | | Hematopoeitic Cell | | |
|---|---|---|---|---|---|---|---|---|---|
| | ABP | Mean ± SD of Enrichment Scores | Percent of Immune-related Terms | ABP | Mean ± SD of Enrichment Scores | Percent of Immune-related Terms | ABP | Mean ± SD of Enrichment Scor es | Percent of Immune-related Terms |
| 1 | **LSP1** | 2.219 ± 0.746 | 37% | **NOD2** | 5.485 ± 3.137 | 33% | **NOD2** | 5.485 ± 3.137 | 33% |
| 2 | **LASP1** | 2.592 ± 0.821 | 13% | **GBP1** | 3.246 ± 1.942 | 46% | **LSP1** | 2.219 ± 0.746 | 37% |
| 3 | **MSN** | 2.617 ± 0.921 | 12% | **GCSAM** | 3.280 ± 1.267 | 39% | **FLII** | 3.348 ± 1.697 | 31% |
| 4 | **CALD1** | 4.733 ± 0.000 | 2% | **ABITRAM** | 5.002 ± 1.713 | 15% | **ABITRAM** | 5.002 ± 1.713 | 15% |
| 5 | **ANXA6** | 2.351 ± 0.607 | 13% | **FLII** | 3.348 ± 1.697 | 31% | **PANX1** | 3.536 ± 1.628 | 24% |
| 6 | **MYO1G** | 1.453 ± 0.137 | 19% | **AIF1** | 3.565 ± 1.895 | 20% | **AIF1** | 3.565 ± 1.895 | 20% |
| 7 | **SCIN** | 2.345 ± 0.000 | 4% | **PANX1** | 3.536 ± 1.628 | 24% | **PSTPIP1** | 2.461 ± 1.141 | 28% |
| 8 | **GSN** | 2.472 ± 0.000 | 2% | **LSP1** | 2.219 ± 0.746 | 37% | **PTK2** | 3.989 ± 2.549 | 7% |
| 9 | **EZR** | 1.945 ± 0.487 | 7% | **PSTPIP1** | 2.461 ± 1.141 | 28% | **CEACAM1** | 2.247 ± 0.437 | 24% |
| 10 | **WIPF1** | 2.116 ± 0.247 | 5% | **PTK2** | 3.989 ± 2.549 | 7% | **VPS16** | 2.817 ± 0.872 | 15% |

**Extended Data Table 2.** Primer sequences for qRT–PCR

| Species | Gene name | Sequences (5’ → 3’) |
|---|---|---|
| *Mus Musculus* | *Lsp1*-Forward | CCAGCCCTTTGGCCTTAGAA |
| | *Lsp1*-Reverse | TGGAAATGGGCAAGGTTGGT |
| | *Mpo*-Forward | AGTTGTGCTGAGCTGTATGGA |
| | *Mpo*-Reverse | CGGCTGCTTGAAGTAAAACAGG |
| | *Cd11b*-Forward | CCATGACCTTCCAAGAGAATGC |
| | *Cd11b*-Reverse | ACCGGCTTGTGCTGTAGTC |
| | *Cd11c*-Forward | CTGGATAGCCTTTCTTCTGCTG |
| | *Cd11c*-Reverse | GCACACTGTGTCCGAACTCA |
| | *Tlr7*-Forward | ATGTGGACACGGAAGAGACAA |
| | *Tlr7*-Reverse | GGTAAGGGTAAGATTGGTGGTG |
| | *Irf7*-Forward | GCGTACCCTGGAAGCATTTC |
| | *Irf7*-Reverse | GCACAGCGGAAGTTGGTCT |
| | *Tbx21*-Forward | AGCAAGGACGGCGAATGTT |
| | *Tbx21*-Reverse | GGGTGGACATATAAGCGGTTC |
| | *Cybb*-Forward | CCTCTACCAAAACCATTCGGAG |
| | *Cybb*-Reverse | CTGTCCACGTACAATTCGTTCA |
| | *C3*-Forward | CCAGCTCCCCATTAGCTCTG |
| | *C3*-Reverse | GCACTTGCCTCTTTAGGAAGTC |
| | *Il1b*-Forward | CCATGACCTTCCAAGAGAATGC |
| | *IL1b*-Reverse | ACCGGCTTGTGCTGTAGTC |
| | *Cebpa*-Forward | CAAAGCCAAGAAGTCGGTGGACAA |
| | *Cebpa*-Reverse | TCATTGTGACTGGTCAACTCCAGC |
| | *Cebpb*-Forward | CAAGCTGAGCGACGAGTACA |
| | *Cebpb*-Reverse | AGCTGCTCCACCTTCTTCTG |
| | *Ets1*-Forward | GATCTCAAGCCGACTCTCACC |
| | *Ets1*-Reverse | GACGTGGGTTTCTGTCCACT |
| | *Blimp1*-Forward | CTTCTCTTGGAAAAACGTGTGGG |
| | *Blimp1*-Reverse | TCATATCAGCGTCCTCCATGT |
| | *Aicda*-Forward | GCCACCTTCGCAACAAGTCT |
| | *Aicda* -Reverse | CCGGGCACAGTCATAGCAC |
| | *Irf4*-Forward | TCCGACAGTGGTTGATCGAC |
| | *Irf4*-Reverse | CCTCACGATTGTAGTCCTGCTT |
| | *Gapdh*-Forward | AGGTCGGTGTGAACGGATTTG |
| | *Gapdh*-Reverse | TGTAGACCATGTAGTTGAGGTCA |
| *Homo* | *LSP1*-Forward | GGAGCACCAGAAATGTCAGCA |

| | | |
|---|---|---|
| *sapiens* | *LSP1*-Reverse | TCGGTCCTGTCGATGAGTTTG |
| | *MPO*-Forward | TGCTGCCCTTTGACAACCTG |
| | *MPO*-Reverse | TGCTCCCGAAGTAAGAGGGT |
| | *IL1B*-Forward | ATGATGGCTTATTACAGTGGCAA |
| | *IL1B*-Reverse | GTCGGAGATTCGTAGCTGGA |
| | *CYBB*-Forward | ACCGGGTTTATGATATTCCACCT |
| | *CYBB*-Reverse | GATTTCGACAGACTGGCAAGA |
| | *ETS1*-Forward | TACACAGGCAGTGGACCAATC |
| | *ETS1*-Reverse | CCCCGCTGTCTTGTGGATG |
| | *GAPDH*-Forward | AAGGTGAAGGTCGGAGTCAA |
| | *GAPDH*-Reverse | AATGAAGGGGTCATTGATGG |

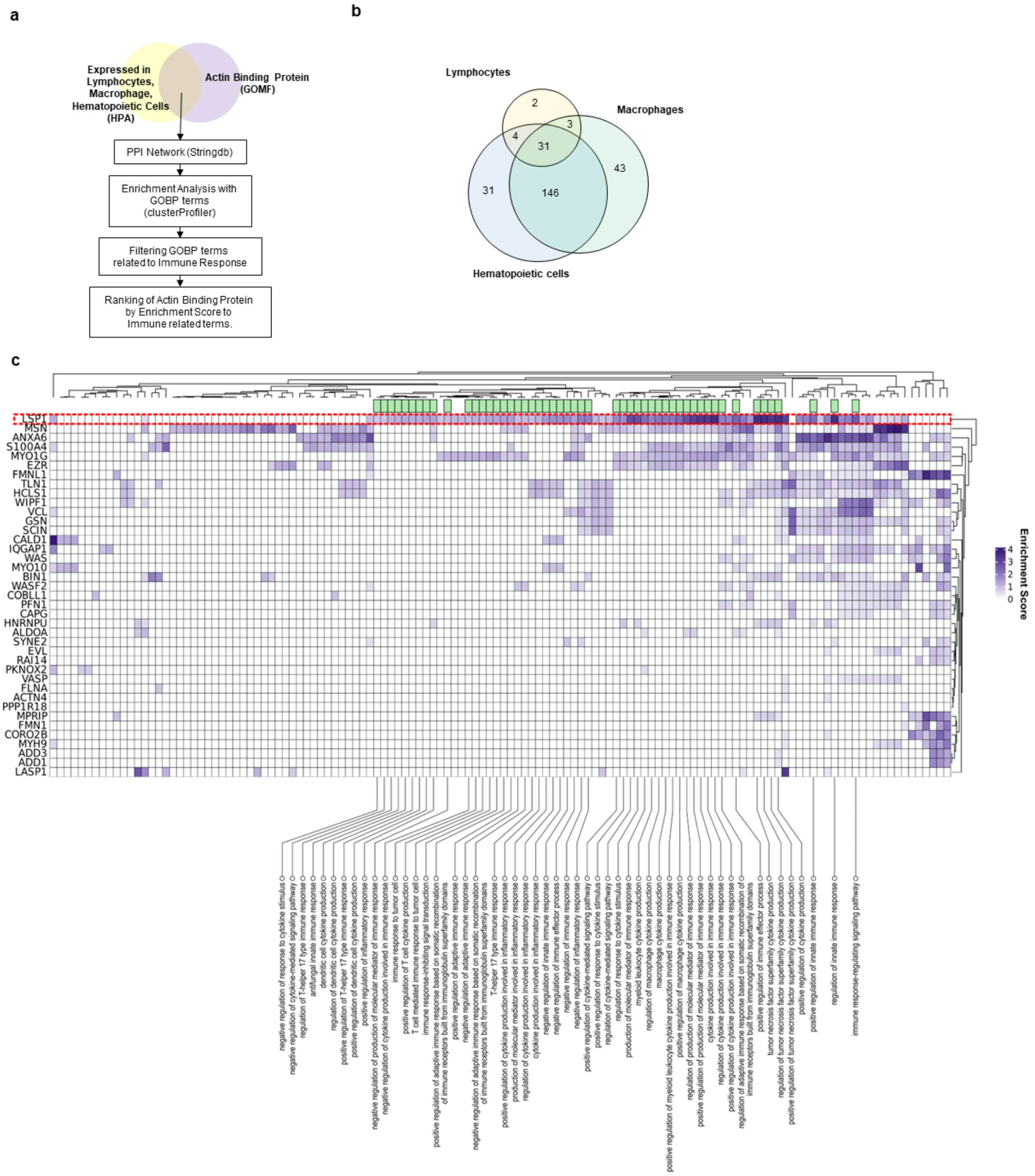


**Extended Data Fig. 1. Identification of actin-binding protein (ABP)s related to immune responses in lymphocytes. a,** Workflow for the analysis of the immunological functions of ABPs expressed in lymphocytes. The Human Protein Atlas (HPA, https://www.proteinatlas.org/), GOMF terms, GOBP terms, and protein–protein interaction (PPI) network were used for the analysis. b, Venn diagram illustrating the counts of ABPs expressed in lymphocytes, hematopoietic cells, and macrophages at the protein level, which were generated based on the HPA database. c, Heatmap displaying the enrichment scores of immune response-related GOBP terms for the 40 ABPs expressed in lymphocytes. The columns highlighted in green at the top indicate the GOBP terms for which LSP1 had a higher enrichment score than did the other 39 ABPs; the terms are described at the bottom.

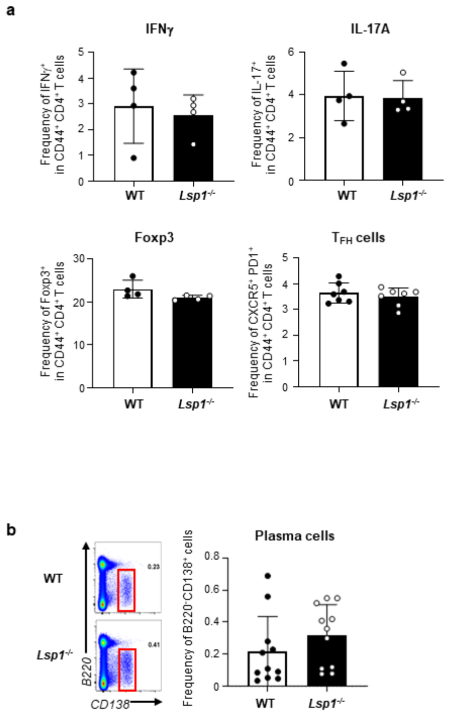


**Extended Data Fig. 2. Effect of LSP1 on the polarization of T helper cells and the generation of plasma cells in the KLH immunization model.** WT and *Lsp1*$^{-/-}$ mice were subcutaneously immunized with 100 μg of KLH emulsified in CFA. On day 7, the draining lymph node cells were harvested and analyzed. **a,** Percentages of IFNγ$^+$, IL-17A$^+$, Foxp3$^+$, and CXCR5$^+$/PD1$^+$ CD4$^+$ T cells in WT versus *Lsp1*$^{-/-}$ mice (n=4 for each group). **b,** Percentages of plasma cells (B220$^-$CD138$^+$ cells; highlighted in red rectangles) in the two groups of mice (n=11 per group). Representative pseudocolor plots are shown on the left. The bar graph represents the percentages of plasma cells in WT and *Lsp1*$^{-/-}$ mice. Data are presented as the mean ± SD of at least two independent experiments. *P* values were determined by the Mann-Whitney *U*-test.

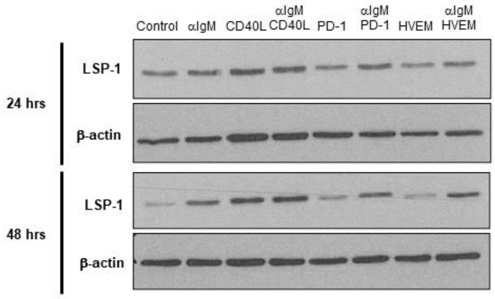


**Extended Data Fig. 3. Upregulation of LSP1 expression in murine B cells by costimulatory signals but not by coinhibitory signals.** Splenic B cells were stimulated with CD40L (0.1 μg/mL), PD-1 (5 μg/mL), or HVEM (5 μg/mL) in the absence or presence of αIgM Ab for 24 or 48 hours. LSP1 expression was then determined via an immunoblot assay. Beta-actin was used as a loading control.

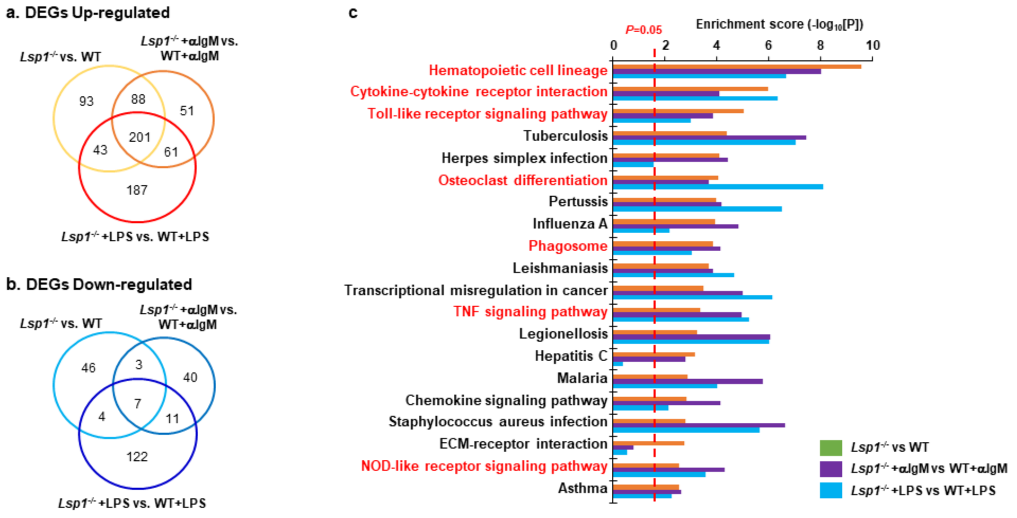


**d. Molecular classes of the genes in Cluster 1 and Cluster 2**

| | Ligand | Receptor | Kinase | TF |
|---|---|---|---|---|
| **Cluster 1** | S100a4, S100a6, Lcp2, Cxcl16, App, Tnf, Fn1, Il1b, Sema4a, Csf3r ,Spp1 ,Ccl5, Tgfbi, Sirpa, Thbs1, Il1f9 ,Lgals3, Socs3, Csf2ra, Lgals1 | Itga5, Cd33, Ms4a3, Cd7, Tlr2, Sema4a ,Csf3r, Clec4e, Ltbr, Siglece, Itgam ,Clec12a, Lair1 ,Csf2ra ,Fcgr3, Pilrb1, Csf2rb, Ms4a4a | Hck, Csf1, Bub1, Fgr, Fes | Atf3, Tfec ,Hmgb2, Cebpa, Cebpb, Ifi204 |
| **Cluster 2** | Il6, Fgl2, Cxcl2, Bcl2a1a, Cish, Il1rn, Il1a, Il15, Insl6 | Itgb7, Fcgr1, Plxna1, Trem1, Fpr1, Itgb2, Ms4a6b ,Cd274, Nrp2, Tnfrsf11a, Procr, P2ry2, Plaur, Cd300e, Hspa1b | Cdk1, Irak3 ,Cdk14, Epha2, Met, Plk2 | Mxd1, Spi1, Parp12, Gfi1b, Tcf19, Klf6 ,Spic, Nfe2l2, Irf1, Plek ,Fosl1 ,Bach1, Kntc1, Jdp2 ,Runx2, Maff, Zfp579, Tceal8 |

**Extended Data Fig. 4. Gene expression profile of splenic B cells from *Lsp1*$^{-/-}$ mice. a, b,** Venn diagram showing the numbers of upregulated (**a**) and downregulated (**b**) DEGs in *Lsp1*-knockout (-/-) B cells stimulated with media alone, αIgM Ab (10 µg/mL), or LPS (2 µg/mL). WT B cells under the same experimental conditions were used as a control. **c,** Bar graph displaying enrichment scores of cellular processes enriched by up- and downregulated DEGs. **d,** Molecular classes of the genes in Cluster 1 and Cluster 2. The DEGs involved in Cluster 1 and Cluster 2 can be categorized into ligands, receptors, kinases, and TFs based on their molecular characteristics.

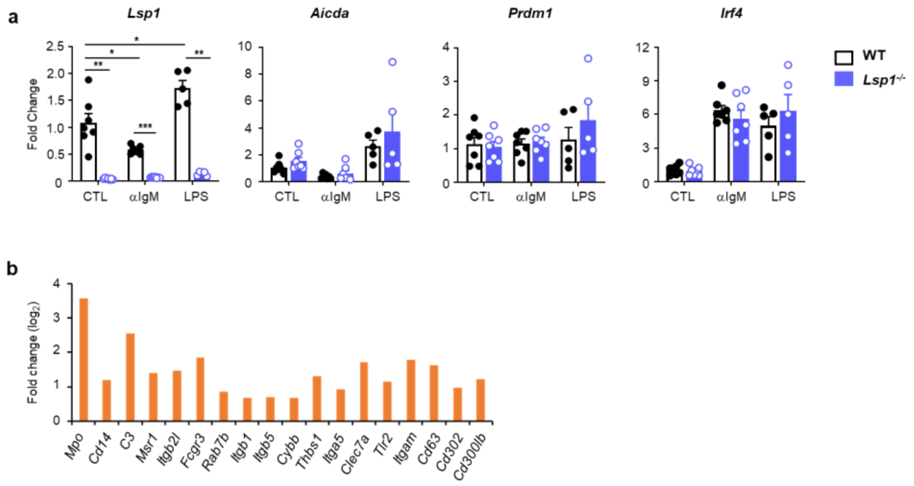


**Extended Data Fig. 5. LSP1 regulation of the genes involved in B-cell differentiation and myeloid cell activation. a,** Quantitative RT–PCR analysis of representative genes related to B-cell differentiation, including *Aicda*, *prdm1*, and *Irf4*, in WT versus *Lsp1*$^{-/-}$ B cells stimulated with media alone, αIgM, or LPS for 4 hours. Fold changes were calculated using the $2^{-\Delta\Delta Ct}$ method. **b,** Bar graph showing fold changes in the expression of phagosome/phagocytosis-associated genes perturbed by *Lsp1* deficiency. The gene sets associated with phagosome/phagocytosis were obtained from Kyoto Encyclopedia of Genes and Genomes (KEGG) pathway database. *P* values were determined by multiple unpaired two-tailed *t* tests (**a**). * $P < 0.05$; ** $P < 0.01$.

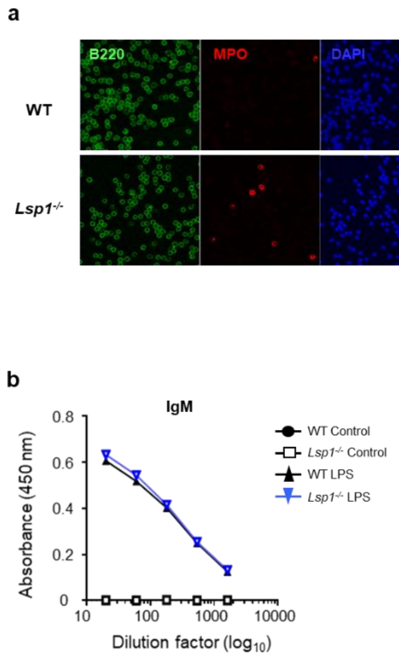


**Extended Data Fig. 6. Effect of *Lsp1* knockout on the frequency of MPO$^{+}$ B cells and the production of IgM. a,** B cells freshly isolated from WT and *Lsp1*$^{-/-}$ mice were fixed and stained with anti-B220 (green) and anti-MPO (red) Abs for immunofluorescence analysis. DAPI was used for nucleic acid staining. Representative images are shown. **b,** IgM production by LPS-stimulated WT versus *Lsp1*$^{-/-}$ B cells (n=3 per group) for 5 days was measured via ELISA.

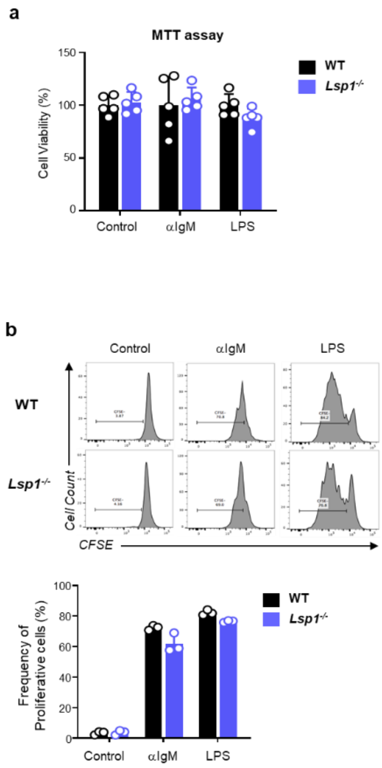


**Extended Data Fig. 7. No effect of *Lsp1* knockout on B-cell survival and proliferation. a,** WT and *Lsp1*$^{-/-}$ B cells were stimulated with αIgM or LPS for 3 days, and cell viability was measured by a 3-[4,5-dimethylthiazol-2-yl]-2,5-diphenyltetrazolium bromide (MTT) assay (n=5 per group). **b,** WT and *Lsp1*$^{-/-}$ B cells were labeled with 0.5 μM carboxyfluorescein succinimidyl ester (CFSE) and were then stimulated as described in (**a**). Representative histograms are shown in the upper panel, and the percentages of proliferative cells are shown in the lower panel. (n=3 per group). Data are presented as the mean ± SD of at least two independent experiments.

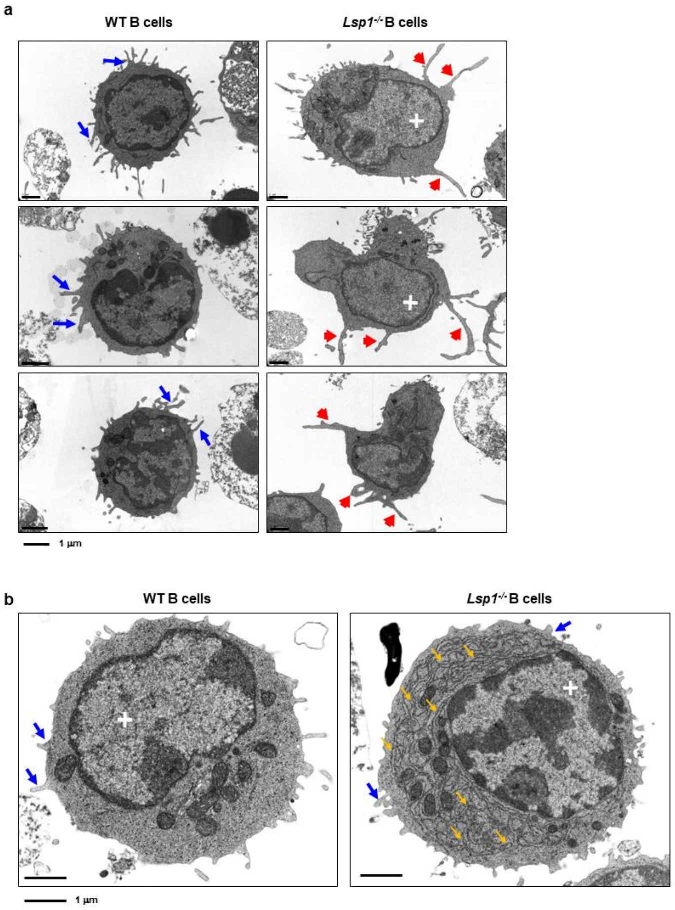


**Extended Data Fig. 8. Ultrastructural analysis of WT and *Lsp1*$^{-/-}$ B cells.** WT and *Lsp1*$^{-/-}$ B cells were stimulated with αIgM (**a**) and LPS (**b**) for 3 days, and their morphologies were analyzed using TEM. When B cells were stimulated with αIgM Abs, they grew in size and exhibited more cytoplasmic organelles, such as mitochondria, endoplasmic reticulum (ER), lysosomes, and extracellular pseudopodia. Overall, the morphology of *Lsp1*$^{-/-}$ B cells was more irregular in shape than that of WT B cells when they were stimulated with αIgM, as indicated by the presence of multiple long pseudopodia (red arrows). In contrast, WT B cells still had a thin cytoplasm with a varying number of small dendrites (blue arrows). Notably, in the nucleus, the proportion of uncondensed chromatin termed 'euchromatin' (white crosses) was increased in *Lsp1*$^{-/-}$ B cells than in WT B cells, which indicates the active transcription of many genes. Upon LPS stimulation, blast formation, which signifies B-cell differentiation to plasmablasts and plasma cells, was frequently observed with increased euchromatin formation (white crosses), mitochondria, and smooth and rough ER in both WT and *Lsp1*$^{-/-}$ B cells. However, compared with WT B cells, *Lsp1*$^{-/-}$ B cells had markedly greater rough ER formation (yellow arrows) spanning from the nuclear envelope to the cytoplasmic membrane. Pseudopodia formation disappeared with LPS stimulation, but the number of small dendrites on the cell surface of *Lsp1*$^{-/-}$ B cells was greater than that on the cell surface of WT B cells. Scale bar, 1 µm.

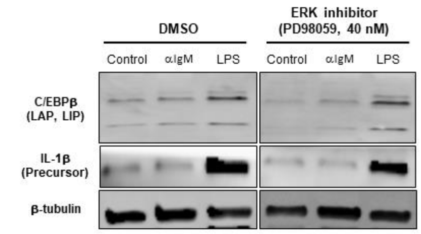


**Extended Data Fig. 9. No change in C/EBPβ expression in *Lsp1*$^{-/-}$ B cells by ERK inhibition.** *Lsp1*$^{-/-}$ B cells were pretreated with ERK inhibitor PD98059 (40 nM) for 1 hour and then stimulated with αIgM or LPS for 24 hours. The expression levels of C/EBPβ and IL-1β (the precursor form) were determined via immunoblot assay. Beta-tubulin was used as a loading control.

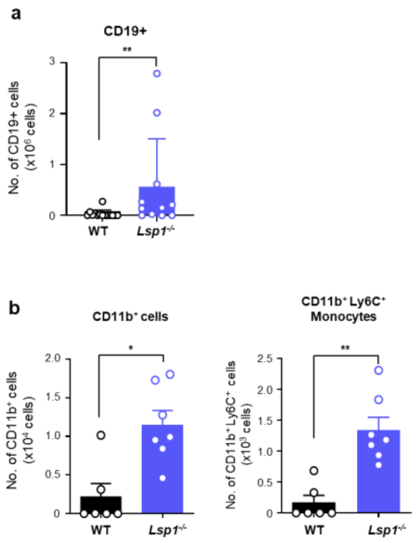


**Extended Data Fig. 10. Comparison of the percentages of B cells and myeloid cells in the BALFs of WT and *Lsp1*$^{-/-}$ mice injected with pristane.** WT and *Lsp1*$^{-/-}$ (n=8 per group) mice were intraperitoneally injected with 500 µl of pristane. After 11 days, the cell numbers of CD19$^{+}$, CD11b$^{+}$, and CD11b$^{+}$Ly6C$^{+}$ cells in the BALF of pristane-injected WT and *Lsp1*$^{-/-}$ mice were determined via flow cytometry. Data are presented as the mean ± SD of at least two independent experiments. *P* values were determined by the Mann Whitney *U* test. * $P < 0.05$; ** $P < 0.01$.

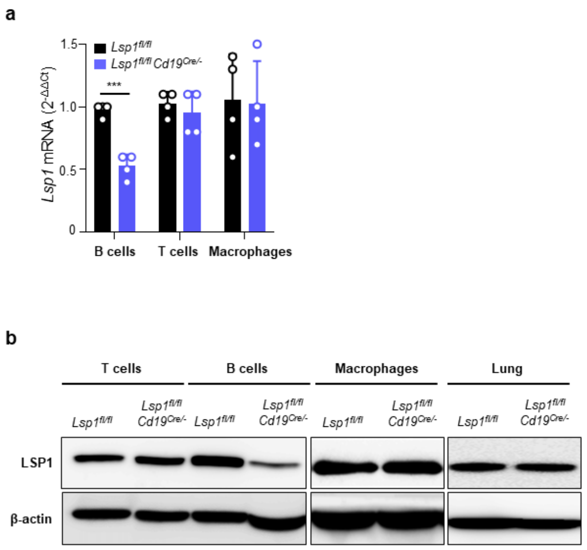


**Extended Data Fig. 11. LSP1 expression in diverse cell types from B-cell-specific *Lsp1*-knockout mice.** LSP1 expression was measured by qRT–PCR (**a**) and immunoblotting (**b**) in the indicated cells and lungs derived from *Lsp1$^{fl/fl}$* and *Lsp1$^{fl/fl}$Cd19$^{Cre/-}$* mice. *P* values were determined by two-way ANOVA with Sidak's multiple comparison test. *** $P < 0.001$.

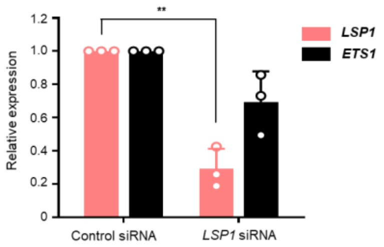


**Extended Data Fig. 12. Effect of *LSP1* deficiency on ETS1 expression in human B cells.** B cells isolated from peripheral blood of healthy donors were transfected with *LSP1* siRNA. After 24 hours, *LSP1* or *ETS1* mRNA expression was assayed via qRT–PCR (n=3). *P* values were determined by two-way ANOVA with Sidak's multiple comparison test. ** $P < 0.01$.

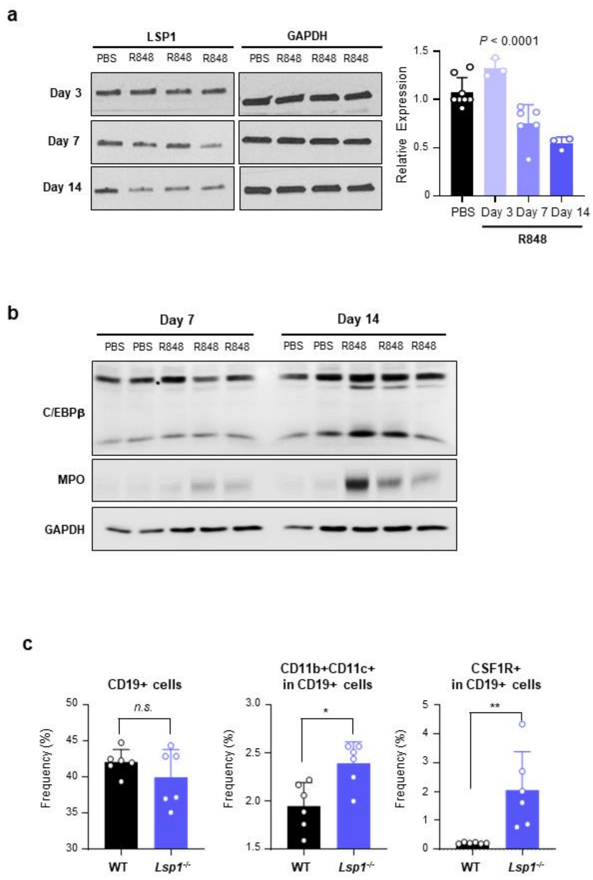


**Extended Data Fig. 13. Changes in the expression of LSP1 and myeloid genes in B cells following repetitive exposure to the TLR7 agonist R848.** WT mice were intraperitoneally injected with 100 µg of R848 every other day, and splenic B cells were isolated on day 3, 7, and 14 after the first injection of R848. **a,** LSP1 expression levels were determined by immunoblotting (left panel) and qRT–PCR (right panel) at each time point. Data in the bar graph represent the mean ± SD of at least two independent experiments. **b,** Expressions of C/EBPβ and MPO were determined via immunoblot assays on day 7 and 14. GAPDH was used as a loading control. **c,** Percentages of total $CD19^+$ B cells, $CD11b^+CD11c^+$ in $CD19^+$ cells, and $CSF1R^+$ in $CD19^+$ cells were determined by flow cytometry 14 days after the first R848 injection. *P* values were determined by one-way ANOVA (**a**) and the Mann Whitney *U* test (**c**). *n.s.*: not significant. * $P < 0.05$; ** $P < 0.01$.

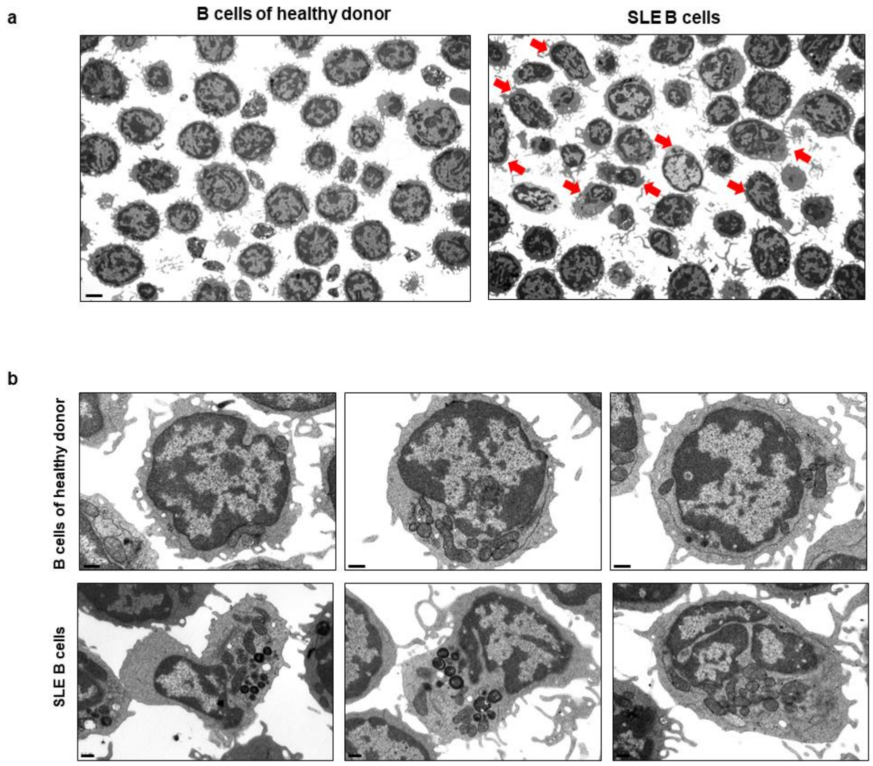


**Extended Data Fig. 14. TEM images of healthy and SLE B cells.** Healthy and SLE B cells were isolated and then fixed to examine the ultrastructure of B cells using TEM. Low- (**a**) and high-magnification images **(b)** are shown.

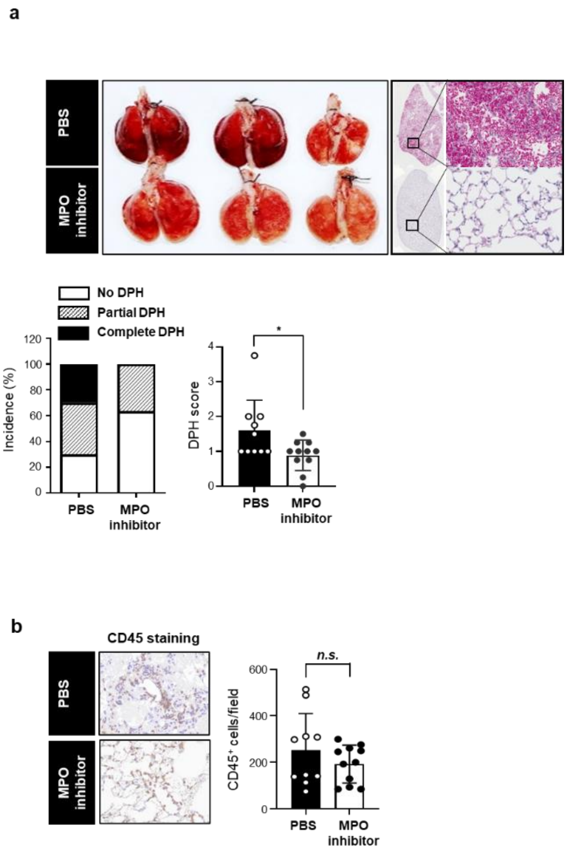


**Extended Data Fig. 15. Amelioration of pristane-induced lupus by an MPO inhibitor.** *Lsp1*$^{-/-}$ mice were orally administered 20 µM of an MPO inhibitor (PF06281355) every other day for 2 weeks in the pristane-induced model. **a,** Representative images of vehicle (PBS, n=10)- or MPO inhibitor (n=11)-treated lung tissues from mice injected with pristane are shown in the top panel. DPH incidence and scores (bottom panel) were determined in the lung tissues as described in **Fig. 5(d)**. **b,** Infiltration of CD45$^{+}$ leukocytes in the lungs of vehicle- versus MPO inhibitor-treated mice was quantified as described in **Fig. 5(e)**. Data are presented as the mean ± SD of at least two independent experiments. *P* values were determined by Fisher's exact test (percentage of DPH incidence in **a**) and the Mann Whitney *U*-test (right panels, **a** and **b**). *n.s.*: not significant. * $P < 0.05$

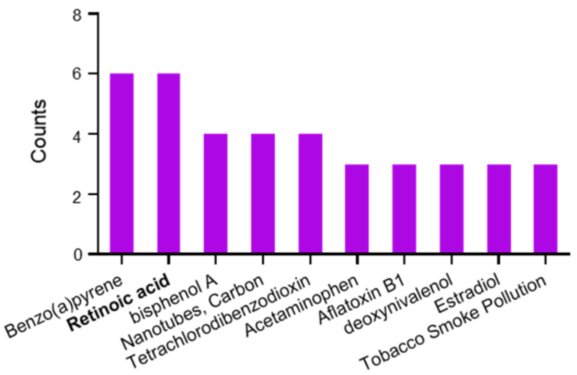


**Extended Data Fig. 16.** The bar graph shows the top 10 compounds that can upregulate LSP1 expression, which were obtained from the Comparative Toxicogenomics Database (CTD, https://ctdbase.org/). The counts on y-axis are the number of references for the chemicals identified as having an interaction with LSP1.

### Extended Data Video Legends

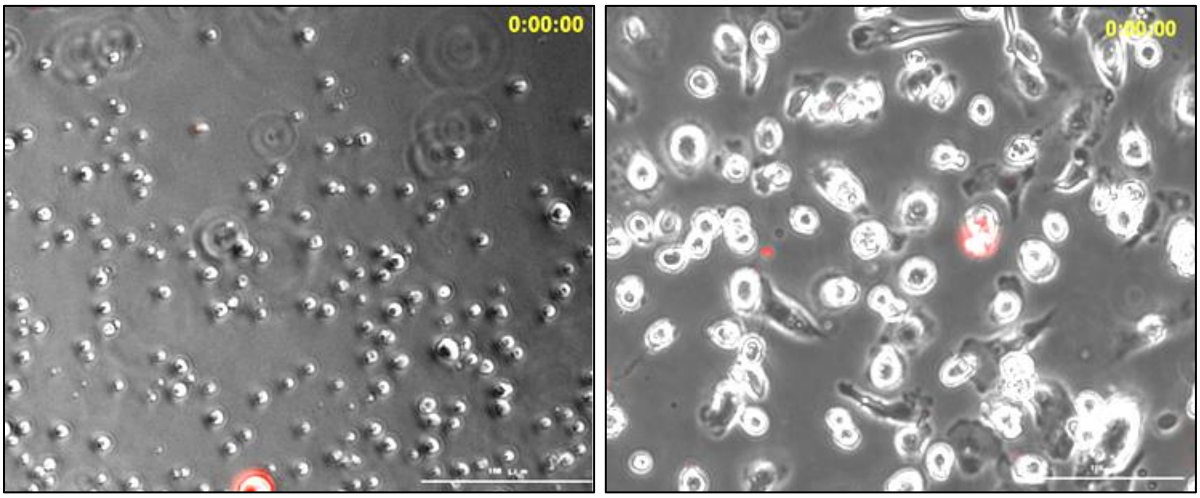


**Extended Data Video 1 and 2. Phagocytic activity of WT (Video 1, left panel) and *Lsp1*$^{-/-}$ B cells (Video 2, right panel).** B cells were stimulated with αIgM for 11 days and treated with Alexa Fluor 594-labeled *E. coli* bioparticles for 30 minutes. Changes in the phagocytic activity and morphology of WT (left panel) and *Lsp1*$^{-/-}$ (right panel) B cells were continuously observed for 30 minutes via LIONHeart live-cell imaging.

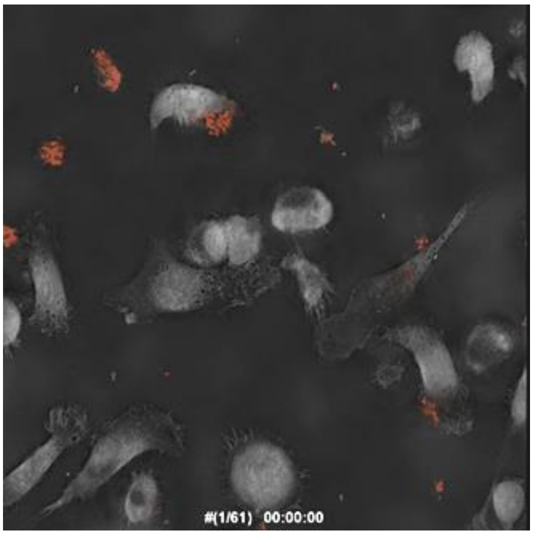


**Extended Data Video 3. TomoCube images showing the phagocytic activity of *Lsp1$^{-/-}$* B cells.** *Lsp1$^{-/-}$* B cells were stimulated with αIgM for 11 days and treated with Alexa Fluor 594-labeled *E. coli* bioparticles for 30 minutes. Changes in the phagocytic activity and morphology of *Lsp1$^{-/-}$* B cells were continuously observed for 30 minutes with a TomoCube Holotomographic microscope.

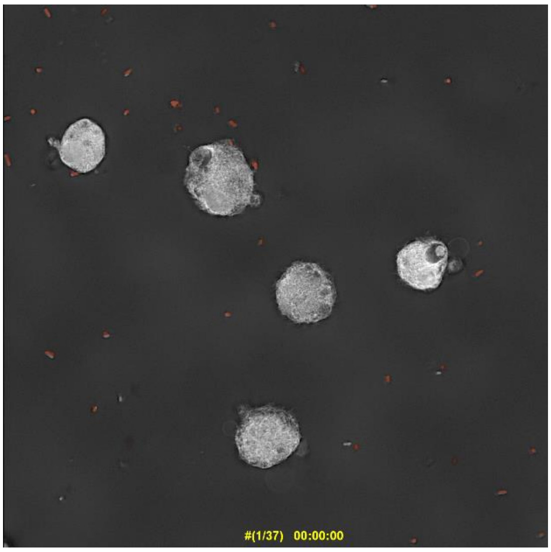


**Extended Data Video 4. Inhibition of the phagocytic activity of *Lsp1*$^{-/-}$ B cells by an actin polymerization inhibitor.** *Lsp1*$^{-/-}$ B cells were stimulated with αIgM for 11 days and then treated with 2.5 μM cytochalasin D, an actin polymerization inhibitor, for 30 minutes, followed by the addition of Alexa Fluor 594-labeled *E. coli* bioparticles for 30 minutes. The phagocytic activity and morphology of *Lsp1*$^{-/-}$ B cells were continuously observed for 30 minutes via a TomoCube Holotomographic microscope.

### Extended Data Methods

#### *Flow cytometry*

Surface staining was performed with the following fluorochrome-conjugated Abs: FITC-CD4 (eBioscience, #11-0041-82), PE/Cy7-CD44 (eBioscience, #25-0441-82), PerCP/Cy5.5-IFNγ (BioLegend, #505821), PE-IL-17A (BioLegend, #506904), Pacific Blue-Foxp3 (BioLegend, #126410), PerCP/Cy5.5-CD4 (BioLegend, #100540), FITC-PD1 (BioLegend, #135214), and APC-CXCR5 (BioLegend, #145506). The stained cells were evaluated with a FACSCanto II (BD Biosciences) instrument with DIVA software. All of the data were analyzed using FlowJo software (FlowJo).

#### *Cell viability and proliferation assay*

The cells were stimulated with αIgM or LPS for 3 days and then treated with (3-(4,5-dimethylthiazol-2-yl)-2,5-diphenyltetrazolium bromide (MTT, 0.5 mg/mL; Roche, #11465007001) for 4 hours. The absorbance at 570 nm was measured by a spectrophotometer (Perkin Elmer), and the data are presented as the percentage of viable cells relative to that of WT B cells. For the cell proliferation assay, B cells were labeled with 1 μM CellTrace™ CFSE (Invitrogen, #C34554) for 10 minutes at room temperature in the dark. The stained cells were washed twice with complete RPMI 1640 media, stimulated with αIgM or LPS for 3 days, and subjected to flow cytometric analysis.

#### *In vivo TLR7 stimulation model*

Mice were intraperitoneally injected with TLR7 agonist R848 (100 μg/mL; InvivoGen,

#tlrl-r848) every other day for 3, 7, or 14 days. The mice were subsequently sacrificed, and splenic B cells were isolated using a mouse pan-B-cell isolation kit II (Miltenyi Biotech, #130-104-443) according to the manufacturer's protocol.
